# Distinct architecture and composition of mouse axonemal radial spoke head revealed by cryo-EM

**DOI:** 10.1101/867192

**Authors:** Wei Zheng, Fan Li, Zhanyu Ding, Hao Liu, Lei Zhu, Jiawei Li, Qi Gao, Zhenglin Fu, Chao Peng, Xiumin Yan, Xueliang Zhu, Yao Cong

## Abstract

The radial spoke (RS) transmits mechanochemical signals from the central pair apparatus (CP) to axonemal dynein arms to coordinate ciliary motility. The RS head, directly contacting with CP, differs dramatically in morphology between protozoan and mammal. Here we show the murine RS head is compositionally distinct from the *Chlamydomonas* one. Our reconstituted murine RS head core complex consists of Rsph1, Rsph3b, Rsph4a, and Rsph9, lacking Rsph6a whose orthologue exists in the *Chlamydomonas* RS head. We present the unprecedented cryo-EM structure of RS head core complex at 4.5-Å resolution and identified the subunit location and their interaction network. In this complex, Rsph3b, Rsph4a, and Rsph9 forms a compact body with Rsph4a serving possibly as an assembly scaffold and Rsph3b in a location that might link the head with stalk. Interestingly, two Rsph1 subunits constitute the two stretching-arms possibly for optimized RS-CP interaction. We also propose a sawtooth model for the RS-CP interaction. Our study suggests that the RS head experiences profound remodeling to probably comply with both structural and functional alterations of the axoneme during evolution.

## Introduction

The radial spoke (RS), consisting of at least 23 proteins, is a conserved T-shaped structure. RS composes of a stalk anchoring on the A-tubule of peripheral doublet microtubules (DMTs) and a head pointing toward the central pair (CP) of MTs (*Luck et al., 1977; Pigino et al., 2011*). It acts as the mechanochemical transducer to transmit signals from the CP of MTs to the dynein arms through mechanical and/or mechanochemical interactions, which regulate the motility of flagellum/cilium (*Brokaw et al., 1982; Goodenough and Heuser, 1985; Huang et al., 1981; Mitchell and Nakatsugawa, 2004; Warner and Satir, 1974; Witman et al., 1978; Yang et al., 2004*). It has been reported that the motility defect in a *Chlamydomonas* CP projection mutant could be rescued by the addition of exogenous protein tags on RS heads, which also suggests the mechanosignaling between CP and RS (*Oda et al., 2014*). Mutants lacking the entire RS complex or part of the RS head will result in paralyzed or abnormal motility of the flagella and cilia (*Sturgess et al., 1979*).

In the RS/CP signal transduction system, RS head is thought to interact with one of the several projections of CP (*Kohno et al., 2011; Wargo and Smith, 2003*). RS head proteins are considered conserved from *Chlamydomonas* (containing RSP1, 4, 6, 9, 10) to mice (containing Rsph1, 4a, 6a, 9, 10b) and human (containing RSPH1, 4A, 6A, 9, 10B) (*Abbasi et al., 2018; Diener et al., 2011; Piperno et al., 1981*). In mammals, mutations in RS head proteins have been linked to primary ciliary dyskinesia (PCD), a genetically heterogeneous recessive disorder of motile cilia that results in neonatal respiratory distress, chronic oto-sinopulmonary disease, male infertility, and organ laterality defects (*Barbato et al., 2009; Bush et al., 2007; Frommer et al., 2015; Knowles et al., 2013; Leigh et al., 2009; Zariwala et al., 2007*). Human patients with mutations in RSPH4A and RSPH9 have PCD due to abnormalities in the RS and loss of the CP MTs in some mutated cilia (*Castleman et al., 2009*). Mouse knockout (KO) models confirmed that Rsph4a is essential for normal ciliary motility and the CP MTs (*Shinohara et al., 2015*). Loss of function mutations in RSPH1 showed a similar phenotype, with abnormal axoneme structures such as defects in the RS and CP of microtubules (*Knowles et al., 2014; Kott et al., 2013*). RSPH6A, homolog of RSPH4A (sharing 67% sequence identity), is specifically expressed in testes and required for sperm flagella formation and male fertility (*Abbasi et al., 2018; Curry et al., 1992; Kott et al., 2013*). Mutations of RSPH3b, the homolog of *Chlamydomonas* stalk protein RSP3, could also contribute to PCD (*Jeanson et al., 2015*). While in *Chlamydomonas*, there is no evidence showing CP loss in the mutant strains of RSP4 and RSP9, indicating that RS may have distinct mechanisms of RS-CP connection in *Chlamydomonas* from that in mammals. Despites the importance of RS and RS head in cilia/ flagella motility, the structural details of RS and the interactions between RS head and CP remains poorly understood, especially in mammals (*Smith and Yang, 2004*).

The general shape of the *in-situ* RS structure in axoneme has been shown by conventional electron microscopy (*Goodenough and Heuser, 1985; Warner, 1970; Warner and Satir, 1974*) and cryo-electron tomography (cryo-ET) (*Bui and Ishikawa, 2013; Knowles et al., 2014; Lin et al., 2014; Nicastro et al., 2006; Pigino et al., 2011*). These studies depicted a T-shaped structure of RS, composed of (1) an elongated stalk that is anchored on the A-microtubule of a peripheral doublet, and (2) an orthogonal head. However, the copy numbers of RS in the 96-nm repeat unit and the shape and size of RS head in different species are different (*Lin et al., 2012; Lin et al., 2014; Pigino et al., 2011*). For instance, it has been reported that there are three RSs (termed RS1, RS2, RS3) in human, sea urchin, and protozoa in the 96-nm repeat; while in *Chlamydomonas* flagella, the density in the RS3 location appears in a much shorter configuration, and lacks structural similarities with spokes RS1 and RS2 (*Bui et al., 2008; Lin et al., 2012*).

The spoke heads of RS1 and RS2 consist of two structurally identical, rotationally symmetric halves that differ from RS3 in the following four species. In human and sea urchin, the RS1 and RS2 spoke heads resemble a pair of ice skate blades (*Lin et al., 2014*); while in the protists and *Chlamydomonas*, the RS heads of RS1 and RS2 have lateral branches that form a connection between the two heads as well as larger interfaces towards the CP. Moreover, the two symmetric halves of RS1 and RS2 appear to have two copies of RS head subunits (*Lin et al., 2012; Lin et al., 2014; Pigino et al., 2011*). Even though the structure revealed by cryo-ET retained biological activity, the resolution is not sufficient to delineate the locations of individual RS subunits. Also, for each of the RS head subunits, there is no available complete high-resolution structure yet. Therefore, a high-resolution structure of RS head complex is necessary to provide a thorough picture of its assembly and how the RS coordinates with CP.

In the present study, our biochemical and functional analyses suggest that in mouse, Rsph6a and Rsph4a may perform comparable functions in different tissues, and Rsph1, Rsph3b, Rsph4a, and Rsph9 can form a stable RS head core complex. We also present an unprecedented cryo-EM structure of the mouse RS head core complex at near-atomic resolution. We unambiguously determined the subunit location and their interaction networks by biochemical studies, electron microscopy analysis of subunit deleted or epitope labeled core complexes, combined with cross-linking and mass spectrometry (XL-MS) analyses. Moreover, based on the fitting of our cryo-EM map of RS head core complex into the available cryo-ET map of human cilia, we propose a sawtooth model in RS-CP interaction. Our results reveal the architecture of the RS head core complex and its subunit interaction network, and provide a model for the mammalian RS-CP interaction, beneficial for our understanding of the mechanism of RS-CP coordination.

## Results

### Mouse Rsph1-Rsph3b-Rsph4a-Rsph9 can form a stable RS head core complex

We first examined whether all the mouse homologous RS head proteins (Rsph1, 4a, 6a, 9 and 10b) and RSP3 (Rsph3b) localize in the axoneme of motile cilia in multiciliated mouse ependymal cells (mEPCs) (*Delgehyr et al., 2015; Zhu et al., 2019*). We found that all these subunits except Rsph6a can be localized in the cilia of mEPCs (Figure 1A). Further sequence comparation showed that Rsph6a is homologous to Rsph4a (with 62.75% sequence identity, (Figure 1—figure supplement 1D). Previous reports have suggested that RSPH6a is obviously enriched in the testis of mammals (*Abbasi et al., 2018*). Consistently, our western blotting analysis demonstrated that Rsph6a was not detected in the cell lysates of mEPCs, but was highly expressed in the sperm (Figure 1—figure supplement 1A). However, distinct from Rsph6a, Rsph4a was absent in the sperm lysates, but enriched in the cell lysates of mEPCs (Figure 1—figure supplement 1A) (*Abbasi et al., 2018*). Furthermore, our immunofluorescent results showed that Rsph6a, but not Rsph4a, was localized at the axoneme of sperm flagella (Figure 1— figure supplement 1B).

**Figure 1.**
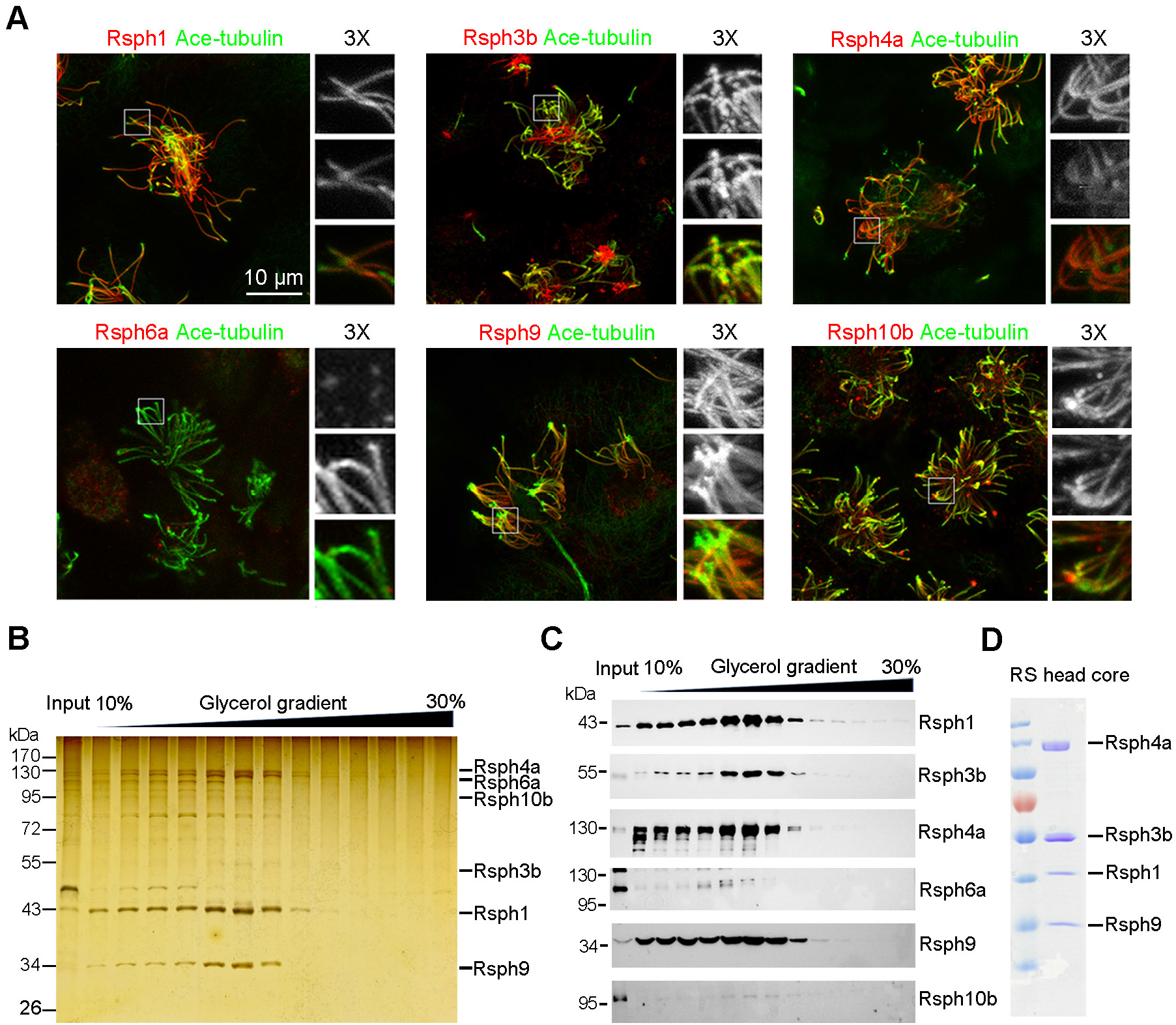
Rsphl-Rsph3b-Rsph4a-Rsph9 can form a stable RS head core complex and be purified. **(A)** Immunofluorescence of mEPCs (mouse ependymal cells) showed that radial spoke head components Rsph1, Rsph3b, Rsph4a, Rsph9, and Rsph10b localize in ciliary axoneme except Rsph6a. Ace-tubulin served as a ciliary axoneme marker. Framed regions were threefold magnified to show details. **(B, C)** Co-expressed RS head complex in HEK293T cells, which was pulled down by flag-tagged Rsph3b followed by performing 10% to 30% glycerol gradient. Silver staining **(B)** and immunoblotting **(C)** results showed Rsph1, Rsph3b, Rsph4a, and Rsph9 co-appear and generate a peak at the seventh fraction, while Rsph6a and Rsph10b can hardly be detected. **(D)** Co-expressed RS head core complex in HEK293T cells and purified using flag beads by flag-tagged Rsph3b and detected by Coomassie-blue staining. Rsph3b carrying a flag tag and Rsph1 carrying a 6×His tag. This purification was repeated more than three times with similar results.

To elucidate the interaction network of the RS head subunits, we performed Co-IP assays, which showed that Rsph4a has strong interactions with Rsph3b, Rsph1, and Rsph9 (Figure 1—figure supplement 1C). Also, we found that Rsph4a and Rsph6a, respectively, can form a stable subcomplex with Rsph1, Rsph9, and Rsph3b (Figure 1—figure supplement 1C). Altogether, our data indicate that in mammals, Rsph6a and Rsph4a may perform comparable functions, and the RS head may contain either Rsph4a or Rsph6a in the motile cilia of mEPCs or the sperm flagella. Therefore, murine RS head is compositionally distinct from the *Chlamydomonas* one which consists of both RSP4 and RSP6.

Furthermore, to examine whether all RS head proteins can form a stable complex *in vitro*, we expressed Flag-tagged Rsph3b together with Rsph1, Rsph4a, Rsph6a, Rsph9, and Rsph10b in 293T cells, and used Flag beads to pull down the complex. The immunocomplex was further subjected to a 10-30% glycerol gradient. It appears that Rsph1, Rsph3b, Rsph4a, and Rsph9 can be purified as a stable complex, whereas Rsph6a and Rsph10 could hardly be detected (Figure 1B, C). We then only expressed the four subunits (Rsph1, Rsph3b, Rsph4a, and Rsph9) that can form the stable RS head core complex for subsequent cryo-EM study (Figure 1D).

### Overall structure of the mouse RS head core complex by cryo-EM

In the sample vitrification process, to overcome the preferred orientation problem associated with the RS head core complex, we used graphene oxide (GO) covered grid with trace amount of detergent (DDM, OG) or polylysine to reduce this effect (Figure 2—figure supplement 1D) (*Ding et al., 2017; Ding et al., 2019; Jin et al., 2019*). We determined the cryo-EM structure of RS head core complex at the nominal resolution of 4.5 Å, which revealed the detailed architecture of the complex for the first time (Figure 2 and Figure 2—figure supplement 1). The map appears to be ~220 Å in height, ~105 Å in length, and ~80 Å in width (Figure 2A-B). This complex has a compact body with two arms stretching out. The two arms are both ~75 Å in height and resembles each other very well, with the lower one (arm 2) slightly better resolved than the upper one (arm 1, Figure 2B). Moreover, visualized from the top, the map appears to have a central canyon dividing the map into two portions (an upper portion and a lower one, Figure 2A), resulting in a pseudo two-fold symmetry of the structure. Interestingly, the front side of the map appears to have a serration-shape with a groove in the middle of this side (Figure 2C).

**Figure 2.**
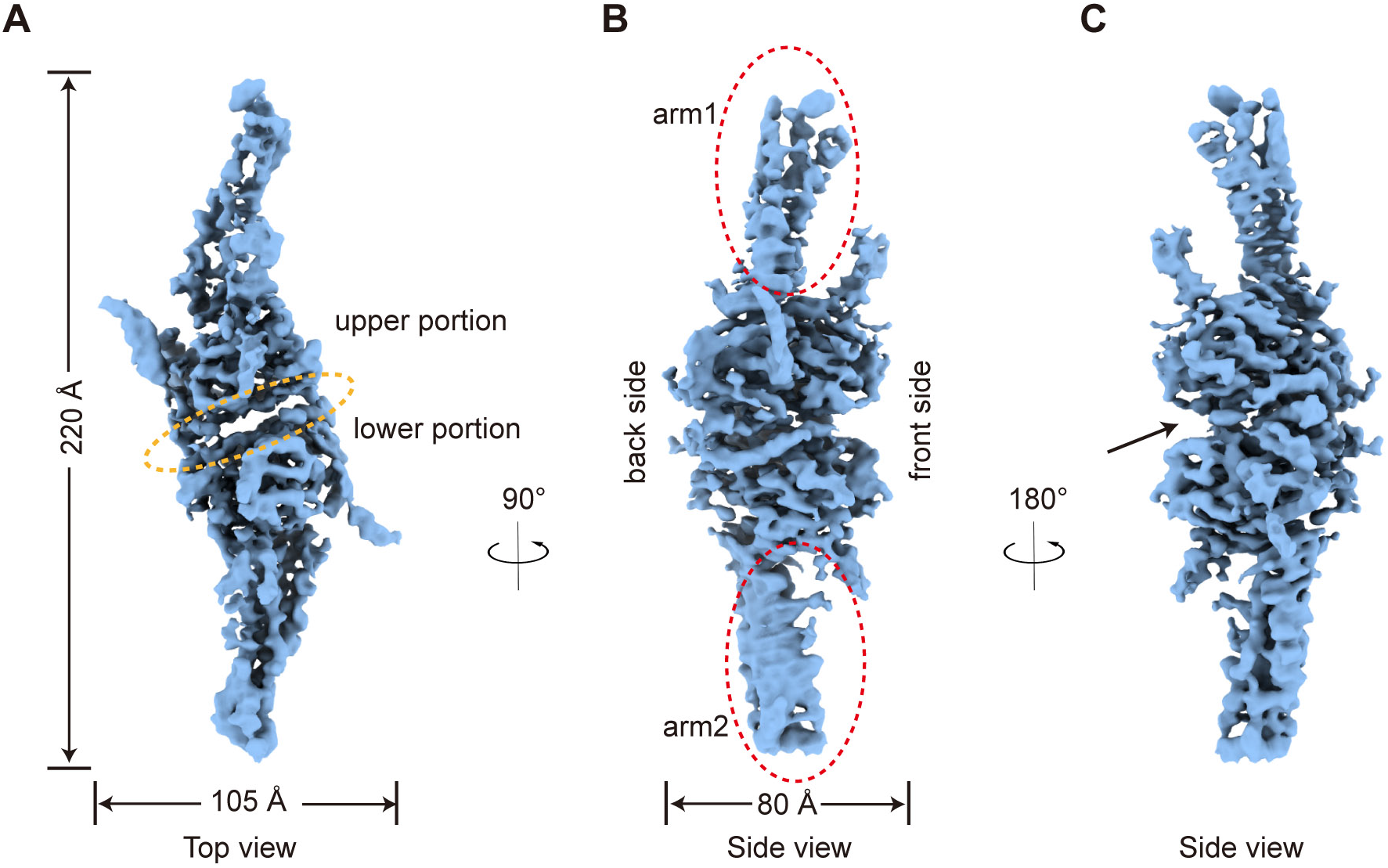
Cryo-EM density map of the RS head core complex. **(A)** Top view of the cryo-EM map of the RS head core complex. There is a central canyon (indicated by an orange ellipsoid) between the upper and lower portions of the map. **(B, C)** Side views of the map, revealing the presence of **(B)** two stretched arms (indicated by red ellipsoid), a body (with front and back side indicated), and **(C)** a groove (indicated by a black arrow) like serration.

### Subunit identification for the RS head core complex by the PA–NZ-1 epitope labeling strategy and XL-MS analysis

To unambiguously locate individual subunit within the complex, we performed cryo-EM analysis on the subunit deleted or epitope labeled RS head core complex based on our previously developed subunit PA–NZ-1 tag-Fab labeling strategy (*Wang et al., 2018a; Wang et al., 2018b*), in combination with XL-MS analysis. To determine the location of Rsph1, we first expressed the complex without Rsph1 (termed RSH^Δ1^). Our reference-free 2D analysis and 3D reconstruction on RSH^Δ1^ both showed that the compacted body remains while the two stretching arms disappeared in comparation to the wild type (WT) core complex (Figure 3A-B, and Figure 3—figure supplement 1A). This result suggests that there are two copies of Rsph1 in the RS head core complex, and each locates in one of the arm position. We then sorted to locate Rsph9 in the map by adopting our PA–NZ-1 epitope labeling strategy. We inserted a dodecapeptide PA tag in the N-terminus of Rsph9 and expressed the PA-labeled core complex (termed RSH^9-PA^). By adding the Fab of the NZ-1 antibody, they can then form a complex, termed RSH^9-PA^–NZ-1. Subsequent reference-free 2D analysis and 3D reconstruction on RSH^9-PA^–NZ-1 both showed an obvious extra density, most likely corresponds to the NZ-1 Fab, exposed outside the lower portion of the body and adjacent to arm2, indicating that Rsph9 may locate in the lower portion of the body close to arm2 (Figure 3C, Figure 3—figure supplement 1B). We then tried to locate Rsph3b in the complex. Inspection of the map depicted a two-long-helix bundle in the lower portion of the body adjacent to the central canyon (Figure 3—figure supplement 2B). Correlatively, sequence analysis by Phyre2 also suggested that, among the four subunits, only Rsph3b contains two extremely long α-helices (~154 aa to 278 aa) (Figure 3—figure supplement 4B, indicated by red underline) (*Kelley et al., 2015*). Thus, Rsph3b might also locate in the lower portion of the body but close to the central canyon. Finally, to locate the remaining Rsph4a subunit, we also followed the PA–NZ-1 epitope labeling strategy. Similar analyses on RSH^4a-PA^–NZ-1 depicted an extra density attached to the back side of the body in the middle upper portion of the map. Considering that Rsph9 and Rsph3b have been located in the lower portion of the body, our analysis suggested that Rsph4a may occupy the upper portion of the body in the complex (Figure 3D, Figure 3—figure supplement 1C).

**Figure 3.**
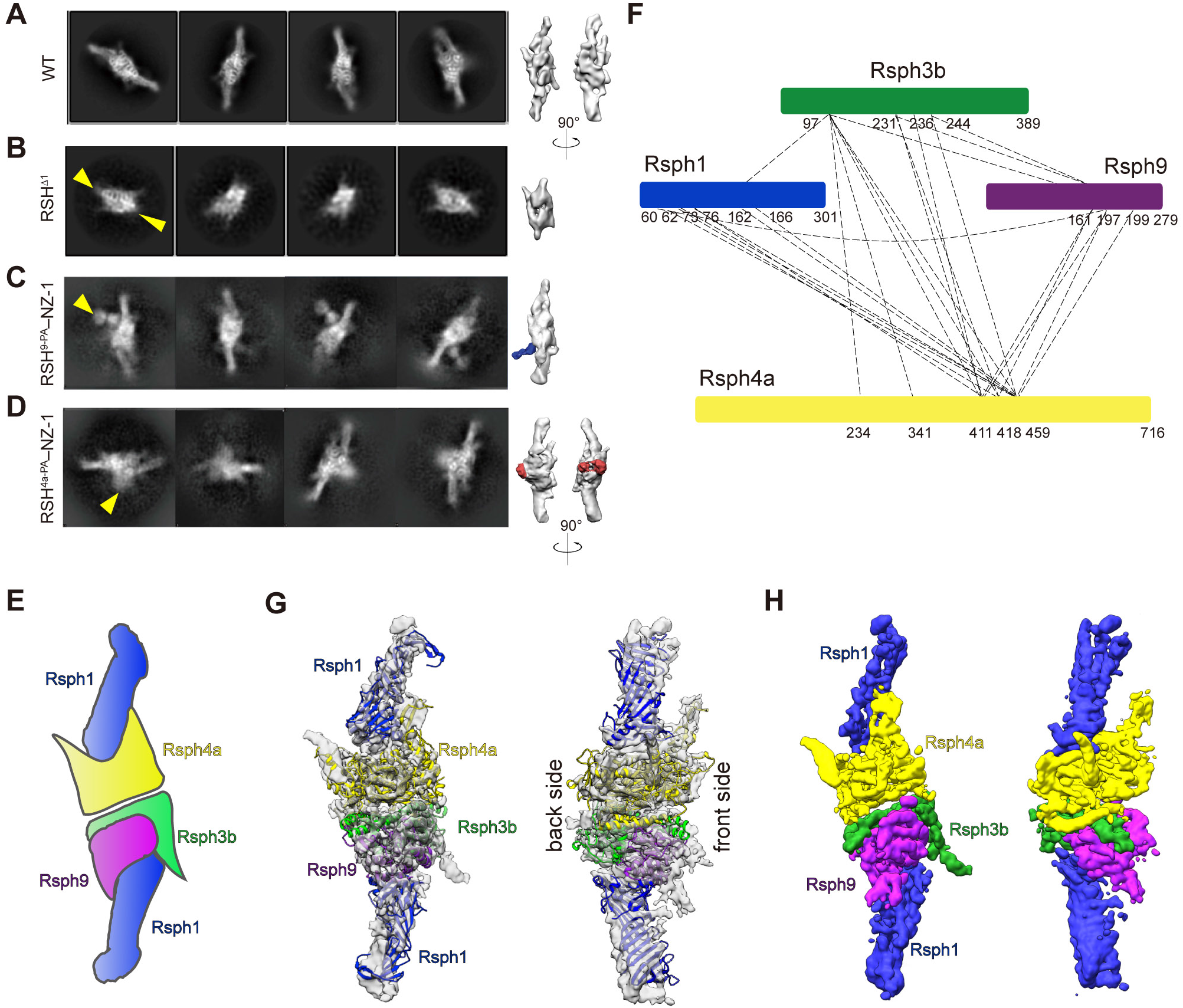
Subunits identification and their interaction network of the RS head core complex. **(A-D)** Determination of subunit locations by cryo-EM analysis on subunit deleted or PA-NZ-1 epitope labeled samples. Compared with the WT RS head core complex **(A)** in both reference-free 2D analysis (left 4 columns) and 3D reconstruction (right 1 or 2 columns), **(B)** RSH^Δ1^ (lacking of Rsph1) retains a compact body but with the two arms missing (indicated by yellow arrow); **(C)** RSH^9-PA^-NZ-1 shows an obvious extra density exposed outside the lower portion of the body and adjacent to arm2 (indicated by a yellow arrow in 2D analysis, and a density in blue in 3D reconstruction), corresponding to the NZ-1 Fab attached to the PA tag inserted to Rsph9; **(D)** RSH^4a-PA^–NZ-1 shows an extra density attached to the back side of the body in the middle upper portion of the map (indicated by yellow arrow in 2D analysis, and a density in red in 3D reconstruction), corresponding to the NZ-1 Fab attached to the PA tag inserted to N-terminus of Rsph4a. **(E)** A cartoon diagram illustrating the potential subunit organization of RS head core complex. Rsph1 in blue, Rsph4a in yellow, Rsph3b in green, and Rsph9 in purple. The color scheme is followed through out. **(F)** XL-MS analysis on RS head core complex, identified cross-linked contacts between different subunits are shown as dotted lines. We used Best e-value (1.00E-02) as the threshold to remove extra XL-MS data with lower confidence. **(G)** Model fitted into the RS head core cryo-EM density map. Here we only fit the Roberta model pieces with high confidence in matching the map SSE features into the density. **(H)** Segmentation of the RS head core cryo-EM map, illustrating the individual subunit location and their interaction network.

Taken together, these structural analyses enabled us to locate the four subunits in the RS head core complex, with two copies of Rsph1 residing in the two arms, Rsph9 in the lower portion of the body close to arm2, Rsph3b also in the lower portion of the body close to the central canyon, and the remaining Rsph4 in the upper core position of the complex (Figure 3E).

Furthermore, to identify the interaction network among the subunits, we performed the XL-MS analysis on the RS head core complex (Figure 3F and Table S2). Our XL-MS data revealed that Rsph4a can be unambiguously cross-linked with all the other three subunits, indicating Rsph4a is likely located in the central position of the complex, which is consistent with our epitope labeling as well as Co-IP assay results (Figure 3D, and Figure 1—figure supplement 1C). Taken together, our data suggest that Rsph4 is located in the core position of the complex, forming strong interactions with all the other subunits (Figure 1—figure supplement 1C). The XL-MS analysis also showed that Rsph3b has multiple cross-links, in the location corresponding to the two long α-helices (~154 aa to 278 aa), with both Rsph4a and Rsph9 (Figure 3F). This is substantiated by the relative central location of Rsph3b determined by our structural analysis (Figure 3E). Moreover, the XL-MS data also revealed multiple interactions between Rsph1 and Rsph4a, in agreement with our structural result (Figure 3E).

### Modeling and subunit interaction network of the RS head core complex

So far, there is no available high-resolution structure for any of the RS head proteins, and consequently template information for homology model building is limited (Figure 3—figure supplement 3A). Also considering the map resolution is not sufficient for de novo model building, the pseudo atomic model building for RS head core complex is challenging. To fulfill this task, we first performed secondary structure element (SSE) prediction by Gorgon based on our cryo-EM map (Figure 3—figure supplement 2A) (*Baker et al., 2011*). We also performed comparative or *ab initial* modeling through Robetta server to predict the structures of the four subunits (Figure 3—figure supplement 3B-E) (*Kim et al., 2004*). Here Gorgon analysis revealed that the two arms appear to consist of a cluster of β-stranded structures (Figure 3—figure supplement 2A). Corroborate to this, sequence analysis and the homology model of Rsph1 both showed this subunit is mostly made up of β-stranded feature (Figure 3—figure supplement 3B, Figure 3—figure supplement 4A). Moreover, our structural analysis has suggested that the two arms are Rsph1 (Figure 3B), we thus fit the Rsph1 model into the two arm regions, which matches the density reasonably well (Figure 3G and Figure 3—figure supplement 2C).

Regarding Rsph9, it has been suggested to locate in the lower portion of the body adjacent to arm1 (Figure 3C, E). Additional SSE prediction indicated a 3-stranded β-sheet also locates in this region, which is supported by the modeling result showing that Rsph9 has a 3-stranded β-sheet feature in the 199-276 aa region (Figure 3—figure supplement 2A, D, and Figure 3—figure supplement 3D). All together, we then placed the Rsph9 model into the lower portion of the body in the map (Figure 3G), which resulted in the reasonable fitting of several small α-helices of Rsph9 into the density (Figure 3—figure supplement 2D). As for Rsph3, our above analyses have located a characteristic 2-helix bundle in the lower portion of the map close to the central canyon, and the model fits in this the corresponding density very well (Figure 3—figure supplement 2B and E). Based on this fitting, the remaining N and C termini of Rsph3b may both reside in the central back side of the map (Figure 3G). Within this complex, the largest subunit Rsph4a however has the least available structural information (Figure 3—figure supplement 3A). Our epitope labeling analysis has located Rsph4a in the upper central portion of the body (Figure 3E), and further SSE analysis suggested the existence of a several stranded β-sheet in this region (Figure 3—figure supplement 2A, indicated by ④). This was substantiated by our modeling result showing Rsph4a has a 6 stranded β-sheet, which fits in the map well with visible β-strand separations (Figure 3—figure supplement 2F).

In this configuration (Figure 3G-H), Rsph1, with a cluster of β-stranded feature, has two copies occupying the two arm regions. The two Rsph1 subunits take up ~60% of the height of the complex, resulting in a stretching-out configuration of the RS head complex, which may serve as a spacer between consecutive RSs and also enlarge the interaction of RS with the apparatus of CP (discussed below). Rsph3b has a two-long-helix bundle passing through the lower central region of the complex. Moreover, Rsph4a is the largest subunit localized in the upper core portion of the complex, interacting with all the other subunits, which may serve as a scaffold for the proper assembly and be important for the function of the RS head core complex.

### The cryo-EM map of RS head core complex matches well with the RS1 and RS2 head envelope density from previous cryo-ET human cilia structure

Within the 96-nm repeat of doublet microtubule (DMT), there are 3 copies of RS complexes (RS1, RS2, and RS3) in the cilia of mammalian systems (*Lin et al., 2014*). We then fit our RS head core cryo-EM map into the head region of the T-shaped RS density from previous cryo-ET map of human cilia (EMD-5950) (Figure 4A) (*Lin et al., 2014*). It appears that the pair of ice-skate-blade-shaped cryo-ET density of the RS head can hold two copies of our RS head core cryo-EM map very well (Figure 4A). This is substantiated by previous cryo-ET studies indicating there might be two copies of RS head subunits in RS1 and RS2 in various systems (*Lin et al., 2012; Pigino et al., 2011*). In our fitting, the RS head core is in an orientation allowing Rsph3b to contact with the stalk of RS. Also considering that RSP3, the ortholog of Rsph3b in *Chlamydomonas*, was regarded as a stalk protein essential for the assembly of the entire RS (*Gaillard et al., 2001; Wirschell et al., 2008*), it is possible that in mammalian RS, Rsph3b may serve as a bridge linking RS head with stalk. Interestingly, although our RS head core structure matches well with the corresponding cryo-ET densities of RS1 and RS2, it doesn’t match the head density of RS3 very well (Figure 4—figure supplement 1). This less well matching to the RS3 head might be due to the following reason: the component of RS3 may not be exactly the same as in RS1 and RS2, related to which, there are only two RSs in *Chlamydomonas* cilia (*Lin et al., 2014; Pigino et al., 2011*). Still, we cannot exclude the possibility that RS3 is rather dynamic and was not fully captured by cryo-ET.

**Figure 4.**
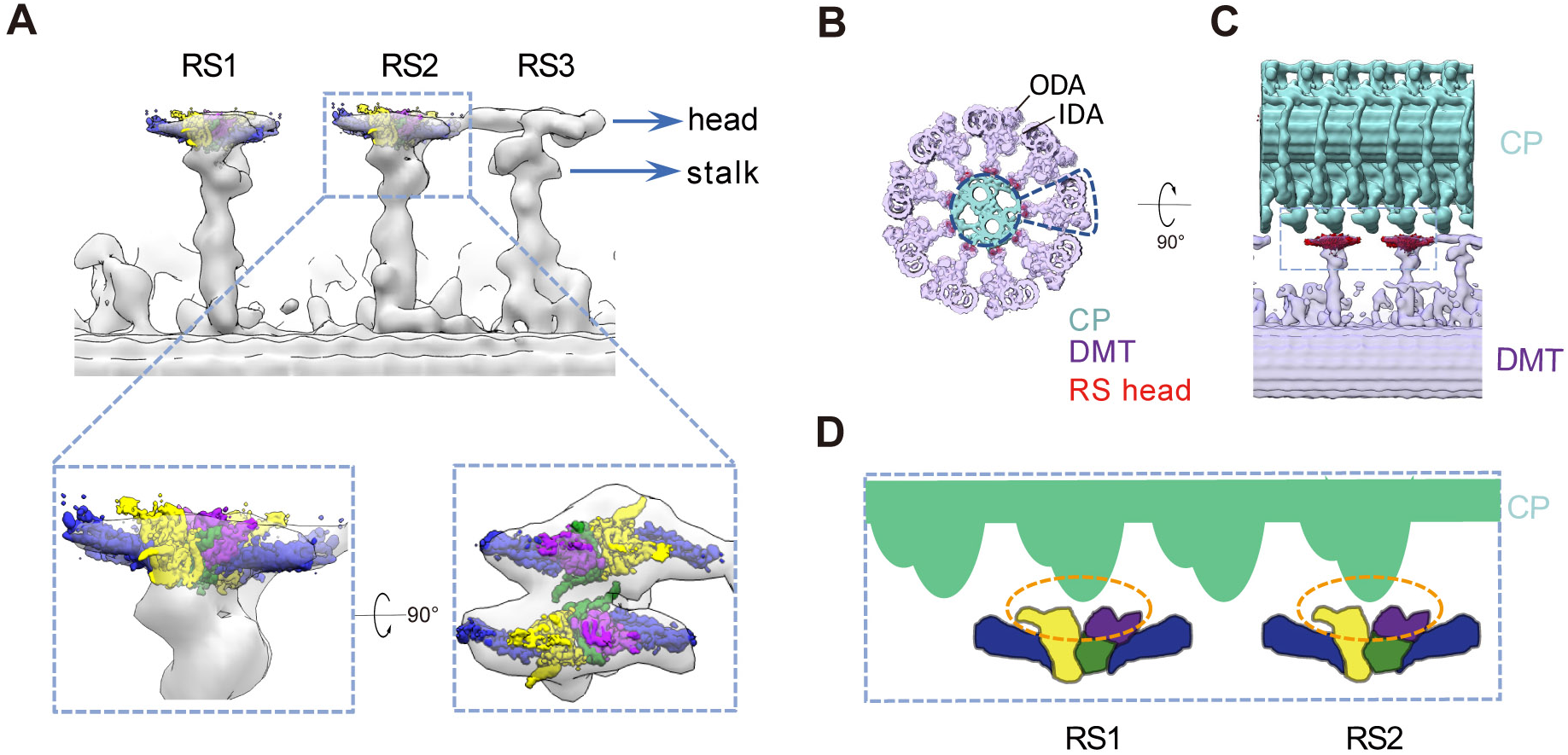
Proposed model of interactions between RS head and central pairs. **(A)** Longitudinal (proximal end of the axoneme to the left) view of RSs from the WT human DMT (transparent grey, EMD-5950) fitted with our cryo-EM map of the RS head core complex (in color, for RS 1 and RS2), and the magnified views of RS2 fitted with our map in different orientations. It appears the ice-skate-blade-shaped RS head density can hold two copies of our RS head core complex very well. In this fitting, the core complex is in an orientation allowing Rsph3b to contact with the stalk of RS. **(B)** Coordinating our RS head core cryo-EM map (in red) into the frame work of DMT-CP based on previous cryo-ET studies on sea urchin CP (EMD-9385, in dark cyan) and human DMT (EMD −5950, in purple). Here the locations of outer dynein arm (ODA) and inner dynein arm (IDA) are indicated. **(C)** Zoom-in view of the CP-DMT interaction network with the RS head core coordinated into CP. **(D)** Based on this fitting, we proposed a sawtooth model in RS-CP interaction (illustrated in the cartoon diagram).

Furthermore, to better visualize the interaction interface between RS and CP, we combined the cryo-ET maps of DMT (EMD-5950, in cooperated with our fitted RS head core complex) with CP (EMD-9385) based on their relative geometry (Figure 4B-C) (*Carbajal-Gonzalez et al., 2013; Ishikawa, 2013*). Based on this fitting, the contacting interface between the RS head core and the CP appears to share a serrated configuration (Figure 4C-D and movie 1). We then speculate that the RS head may rely on the serrated contacts to stabilize CP and to receive signals from CP for further signal transduction through RS head to RS stalk and the inner dynein.

## Discussion

The radial spoke head, the distal of RS close to CP, plays vital roles in the RS-CP interaction. Here, our biochemical and functional results suggest that Rsph6a and Rsph4a may perform comparable functions in different tissues, and mouse RS head is compositionally distinct from the *Chlamydomonas* one. Mouse RS head core complex consists of Rsph1, Rsph3b, Rsph4a, and Rsph9, lacking Rsph6a whose orthologue exists in the *Chlamydomonas* RS head. Consistently, Rsph6a was not expressed in multiciliated murine ependymal cells. We also showed that Rsph3b, usually considered as a stalk protein in *Chlamydomonas*, is a component in the mouse RS head complex. We resolved the cryo-EM structure of the RS head core complex revealing the unforeseen architecture of the complex. Through subunit deletion and PA-NZ-1 epitope labeling strategies, we located the subunits within the complex, and further delineated the subunit interaction network by XL-MS, Co-IP, and structural modeling analyses. In this configuration: (1) two copies of Rsph1 stretch out forming two arms that may enlarge the interaction between RS and CP apparatus; (2) Rsph3b, Rsph4a, and Rsph9 form the compact body, with Rsph4a interacting with all the other subunits and may serve as a scaffold for the complex assembly; (3) Rsph3b is in a location which may bridge the RS head with the stalk. Our study reveals the complete architecture and subunit locations of the RS head core complex, and enabled us to propose a sawtooth model in RS-CP interaction, which together facilitates our understanding on the mechanism of RS-CP interaction and the role of RS head in cilia motility.

### The homologous Rsph4a and Rsph6a are tissue specific components in RS head

Here we showed that the expression of the homologous Rsph4a and Rsph6a is tissue specific (Figure 1A and Figure 1—figure supplement 1A-B). This suggests that RS head may have different compositions in distinct tissues, which is substantiated by previous report showing that *Rsph6a* is testis-specific in human and mice (*Abbasi et al., 2018; Kott et al., 2013*). The motility patterns of cilia in mammalian tissues are distinct, e.g. it adopts planar beating pattern in the airway, brain and oviduct multiciliated cells, whip-like beating pattern in male efferent duct multiciliated cells, undulatory beating pattern in matured sperm, and rotation pattern of node cilia in mouse embryo (*Brokaw, 2009; Hirokawa et al., 2006; Satir et al., 2014; Shinohara et al., 2015*). We thus propose that the motility pattern difference may be to some extend related to the compositional differences of RS in different tissues, i.e. the nodal cilia lacking of RS shows a rotational moving pattern (*Hirokawa et al., 2006*), the ependymal cells in brain lacking of Rsph6a shows a planner beating pattern, the matured sperm lacking of Rsph4a shows an undulatory beating pattern. (*Abbasi et al., 2018*). Future structural study of RS head containing Rsph6a could help to understand the mechanism of different motility pattern of cilia and flagella. In addition, previous study demonstrated that the size of RS head is dramatically distinct between mammalian cilia and protozoan (*Lin et al., 2014*). The different size of RS head may play diverse mechanical contacting role in RS-CP interaction, which indicates that cilia in different species may use varied compositions of RS head subunits to form dissimilar shape to function in RS-CP interactions.

### Rsph1 may play an important role in enlarging the RS-CP interaction

It has been documented that deletion of any of the four subunits (Rsph1, Rsph3b, Rsph4a, Rsph9) of the core complex is related to PCD diseases, indicating all these subunits may be involved in direct or indirect interactions with CP for receiving signals from CP (*Castleman et al., 2009; Jeanson et al., 2015; Kott et al., 2013*). Our structural study showed that Rsph3b, Rsph9, and Rsph4a form a compact body. Deletion of any of these subunits may affect the assembly of the body and the attachment of two copies of Rsph1 to the body, and eventually disrupt the proper assembly of RS head.

Moreover, our structure suggests that the two Rsph1 arms occupy ~60% of the height of the complex, which may act as a spacer between neighboring RSs and also greatly enlarge the interaction between RS and the central pair apparatus. Based on our current cryo-EM structure as well as the previous cryo-ET structures (*Lin et al., 2014; Pigino et al., 2011*), the enlarged sheet-like RS head may act as the pad of the bicycle breaks, that can form close contact with the CP apparatus, and together with the stalk to support the CP. We also showed that when the two Rsph1 arms are deleted, the remaining body of RS head appears smaller (Figure 3B), which might affect the proper interaction between RS and CP apparatus and also with the neighboring RS3, which together may eventually lead to PCD disease (*Knowles et al., 2014*). Correlatively, the RS heads of RS1 and RS2 appear invisible in the cryo-ET map of human cilia with deleted RSPH1 (*Lin et al., 2014*), which may be due to the reduced size of RS head and thus reduced interaction with CP apparatus and the neighboring RS, the reaming compact body attached with stalk become dynamic, and hardly be resolved through cryo-ET analysis. Consequently, the CP could have various abnormalities, even completely disappeared (*Lin et al., 2014*). Altogether, Rsph1 may play a role in enlarging the interaction between RS and CP.

Also in that study in human cilia, the head of RS3 remains although that of RS1 and RS2 disappeared when RSPH1 was deleted (*Lin et al., 2014*). This indicates that, compared with RS1 and RS2, the head of RS3 may have a distinct composition without Rsph1, and it is possible that Rsph10b, sharing the common MORN motif with Rsph1, may replace Rsph1 in RS3 (*Yang et al., 2006*).

### Rsph3b is coordinated into the RS head and may connect the RS head with stalk

Previous study suggested that in *Chlamydomonas*, RSP3 subunit (homologous to Rsph3b) is a stalk protein located near DMT, and RSP3 may form a dimer essential for the assembly of the entire RS (*Gaillard et al., 2001; Jivan et al., 2009; Wirschell et al., 2008*). Here we showed that the mouse Rsph3b is coordinated into the RS head and locates in the core region of the head bridging the head with stalk (Figure 3H), and Rsph3b is considerably shorter than that of *Chlamydomonas* RSP3 (Figure 4—figure supplement 2). Taken together, in different species, the same subunit may occupy distinct locations within the complex and hence play different roles in the assembly or function of the complex. Still, how many copies of Rsph3b in the complete RS complex and how Rsph3b interacts with other RS stalk proteins remain to be elucidated.

### Mechanism of RS-CP interactions

Based on the fitting of our cryo-EM map into the framework of DMT-CP density of previous cryo-ET studies (*Carbajal-Gonzalez et al., 2013; Lin et al., 2014*), the serration-shaped front side of the RS head appears to face the CP projections (Figure 4). Interestingly, we also observed the serration-shaped feature in the CP projections from the previous cryo-ET result (*Carbajal-Gonzalez et al., 2013*). Altogether, we propose a sawtooth mechanism in RS-CP interaction, that is RS head may use the serrated contacts formed between RS head and CP projections to stabilize the RS-CP interaction and receive signals from CP, which could lead to cilia motility by cooperating with dynein arms (Figure 4B-D).

## Materials and Methods

### Protein expression and purification

The full-length cDNAs for Rsph1 (NM_001364916.1), Rsph3b (NM_001083945.1), Rsph4a (NM_001162957.1), Rsph6a (NM_001159671.1), Rsph9 (NM_029338.4), and Rsph10b (XM_006504895) were amplified by PCR from total cDNAs from mouse testis or murine tracheal epithelial cells (MTECs) and constructed into pMlink-His, pMlink-HA, pcDNA3.1-NFLAG to express FLAG tagged Rsph3b, His tagged Rsph1, HA tagged Rsph6a and no tagged Rsph4a and Rsph10b.

When the HEK293T cells density reached 80-90% confluency, the cells were transiently transfected with the expression plasmids and polyethylenimines (PEIs) (Polysciences). For expressions of the RS head component, the plasmids for each of the individual RS head subunit were premixed in equal proportion leading to a total plasmid of 20 μg, then mixed with 60 μg of PEIs in 1 ml of OPTI-MEM for 20 mins before application. For transfection, 50 ml of the mixture was added to 50 dishes of cell culture and then replaced medium after 8 hrs. Transfected cells were cultured for 48 hrs before harvest. For purification of the RS head core complex, the HEK293T cells were collected and resuspended in the lysis buffer containing 25 mM Tris pH 8.0, 150 mM KCl, 0.1% NP-40, 1 mM EDTA, 10 mM Na_4_P_2_O_7_, 10% Glycerol, and protease inhibitor cocktails (Calbiochem, 539134, Billerica). After sonication on ice and centrifugation at 20,000g for 1 hr, the supernatant was collected and applied to anti-Flag M2 affinity resin (Sigma). The resin was rinsed with wash buffer containing 20 mM HEPES pH 7.5, 150 mM KCl, 10% Glyceol, and protease inhibitors. The protein was eluted with wash buffer plus FLAG peptide (1 mg/ml).

The eluent was then crosslinked with 0.1% glutaraldehyde (Sigma) for 3 hrs and neutralized by adding glycine (pH 7.5) to a final concentration of 50 mM for 1 hr at 4°C. After concentrated to 0.5 mL, the protein was further purified by gradient centrifugation using a 10–30% (w/v) glycerol gradient. The resulting gradients were subjected to ultracentrifugation at 4°C for 14 hrs at 41,000 rpm in an SW41 Ti rotor (Beckman), and then fractionated. The fractions were concentrated for biochemical and cryo-EM analyses.

### Cell culture and immunofluorescent

Cells were maintained at 37°C in an atmosphere containing 5% CO_2_. Unless otherwise indicated, the culture medium was Dulbecco’s Modified Eagle’s medium (DMEM) supplemented with 10% fetal bovine serum (Biochrom, Cambridge, UK), 0.3 mg/ml glutamine (Sigma), 100 U/ml penicillin (Invitrogen), and 100 U/ml streptomycin (Invitrogen). Multiciliated mouse ependymal cells were obtained and cultured as described (*Zhu et al., 2019*). Briefly, the telencephalon taken from the P0 C57BL/6J mice were digested and spread onto laminin coated flasks. After neurons were shaken off and removed, the cells were transferred to laminin coated 29mm glass bottom dishes (Cellvis, D29-14-1.5-N) and then starved in serum free medium to induce differentiation into ependymal cells.

For immunofluorescent, spermatozoa collected form the 6 weeks C57BL/6J mice were diluted in PBS, then spread onto laminin-coated 29-mm glass-bottom dishes (Cellvis, D29-14-1.5-N). Then Spermatozoa and mEPCs were pre-extracted with 0.5% Triton X-100 in PBS for 30 secs, followed by fixation with 4% paraformaldehyde in PBS for 10 min at room temperature and permeabilization with 0.5% Triton X-100 for 15 min. Immunofluorescent staining was carried out as described (*Zhao et al., 2013*). Antibodies used are listed in Table S3.

Confocal images were captured by using Leica TCS SP8 system with a 63×/1.40 oil immersion objective and processed with maximum intensity projections. A 592-nm depletion laser was used for Alexa Fluor 488 dye. A 660-nm depletion laser was used for Alexa Fluor 546, −555, and −594 dyes. Images were processed by Leica LAS X software.

### Co Immunoprecipitation (Co-IP) and Western blot

Co-IP was performed as described previously (*Zhao et al., 2013*). Briefly, HEK293T cells transfected for 48 hr were lysed with the lysis buffer [20 mM Tris-HCl (pH 7.5), 150 mM KCl, 0.5% NP-40, 1 mM EDTA, 10 mM Na4P2O7, 10% Glycerol]. Pre-cleared cell lysates were incubated with anti-FLAG beads (Sigma, A2220) or anti-HA beads (Sigma, E6779) for 3 hrs at 4°C. After three times of wash with the lysis buffer and wash buffer [20 mM Tris-HCl (pH 7.5), 150 mM KCl, 0.5% NP-40, 1 mM EDTA, 10 mM Na_4_P_2_O_7_, 10% Glycerol], proteins were eluted with 30 μl of 1 mg/ml FLAG peptide or HA peptide.

For western blot, proteins separated by SDS-PAGE were transferred to nitrocellulose membranes (General Electric Company). Blots were pre-blocked with 5% non-fat milk diluted in TBST (50 mM Tris-HCl, 150 mM NaCl, 0.05% Tween-20, pH 7.5) for 1 hr and then incubated with primary antibodies. After washing with TBST for four times, membranes were incubated with secondary IgG-HRP antibodies. After three times wash in TBST, protein bands were visualized with enhanced chemiluminescent reagent (PerkinElmer) and exposed to X-ray films (Carestream) or Mini Chemiluminescent Imager (MiniChemi 610 Plus, Beijing Sage Creation Science Co).

### Cryo-EM sample preparation and data collection

Freshly purified RS head core complex (0.4 mg/ml) was placed onto glow-discharged grids (Quantifoil R1.2/1.3 200 mesh Cu grids) and plunge-frozen into liquid ethane, cooled with liquid nitrogen, using a Vitrobot Mark IV (FEI, now Thermo Fisher Scientific). The vitrified RS head core complex showed an obvious preferred problem. To overcome this problem, we tried several vitrification conditions, including using graphene oxide supports, grid coated with polylysine (*Ding et al., 2019; Jin et al., 2019; Zang et al., 2016*), or sample with added detergent OG or DDM, which allowed a slightly improved broader orientation distribution of the complex. For PA inserted RS head core complex, the sample was incubated with NZ-1 Fab in a molar ratio of 1:2 (RS head core vs. NZ-1 Fab) on ice for 30 min, and then prepared for vitrification as described.

Selected grids were imaged in a Titan Krios transmission electron microscope (FEI, now ThermoFisher Scientific) operated at 300 kV at liquid nitrogen temperature. A total of ~5,500 movies were collected with a nominal magnification of 22,500×. The images were recorded on a Gatan K2 Summit direct electron detector operated in super resolution mode, yielding a pixel size of 1.02 Å after 2 times binning. Each frame was exposed for 0.2 s, with an accumulation time of 7.6 s for each movie, thus leading to a total accumulated dose of 58 e^-^/Å^2^ on the specimen.

### Image processing

We used command line MotionCor2 (*Zheng et al., 2017*) to align the 38 frames to obtain a single micrograph with dose weighted, and CTFFIND4 (*Rohou and Grigorieff, 2015*) in Relion3 (*Zivanov et al., 2018*) to determine the contrast transfer functions. Particles were picked in EMAN2 (*Tang et al., 2007*) and imported into Relion3 for 2D classification. To reduce the preferred-orientation problem, we performed another round of 2D classification on the preferred-particles. After removing bad particles and also randomly reducing the preferred-particles, we combine them with the particles in other views. 740,463 particles were used for further multiple rounds of 3D classification, with the initial model generated by utilizing EMAN1 (*Ludtke et al., 1999*). 186,398 good particles remained for 3D auto-refinement in Relion3, which were further refined in cisTEM (*Grant et al., 2018*), and sharpened by applying a negative B-factor of −90 Å^2^. The overall resolution of 4.5 Å was estimated based on the gold-standard criterion using a FSC of 0.143 C (*Rosenthal and Henderson, 2003; van Heel and Schatz, 2005*). The local resolution was estimated by using ResMap (*Kucukelbir et al., 2014*).

### Model building

As described in the text, the available structural information for RS head proteins are very limited, also the resolution of the map is not sufficient for de novo model building. We first utilized Gorgon (*Baker et al., 2011*), an interactive molecular modeling system specifically geared towards cryo-EM density maps of macromolecular complexes, to build a density skeleton, which is a compact geometrical representation of the density map using curves and surfaces (*Baker et al., 2007*). Combining the skeleton and SSEs determined against the density map by Gorgon, with our results on the subunit locations within the map and the predicted structures by Robetta server (*Kim et al., 2004*), we were enabled to place most the structural models into the map (Figure 3G-H). Several obvious features in the density match the corresponding models reasonably well, including a cluster of β-strand in Rsph1, several small α-helices in Rsph9, the two long α-helices in Rsph3b, and the six-turn-β-strand in Rsph4 (Figure 3—figure supplement 2C-F).

### Cross-linking mass spectrometry

The purified RS head core complex from glycerol gradient was cross-linked by Bis[sulfosuccinimidyl] (BS3), with a final concentration of crosslinker at 2.5 mM on ice for 4 hours or RT for 1 hour. 50 mM Tris-HCl was used to terminate the reaction after incubation. Cross-linked complexes were precipitated with cooled acetone and lyophilized. The pellet was dissolved in 8 M urea, 100 mM Tris pH 8.5, followed by TCEP reduction, iodoacetamide alkylation, and overnight trypsin (Promega) digestion. Digestion was quenched by 5% formic acid. Tryptic peptides were desalted with MonoSpin C18 spin column (GL Science) and then separated within a home packed C18 column (Aqua 3 cm, 75 cm × 15 cm, Phenomenex) in a Thermo EASY-nLC1200 liquid chromatography system by applying a 60-minute step-wise gradient of 5–100% buffer B (84% acetonitrile (ACN) in 0.1% formic acid). Peptides eluted from the LC column were directly electrosprayed into the mass spectrometer with a distal 2 kV spray voltage. Data-dependent tandem mass spectrometry (MS/ MS) analysis was performed with a Q Exactive mass spectrometer (Thermo Fisher, San Jose, CA). Raw data was processed with Plink software (*Yang et al., 2012*) and Proteome Discoverer 2.2 xlinkx (Table S2).

## Acknowledgements

We thank Z. Zhou and L. Bao (Institute of Biochemistry and Cell Biology) for their suggestions and supports on this project. We are grateful to the staff of the NCPSS Electron Microscopy facility, Database and Computing facility, Mass Spectrometry facility, and Protein Expression and Purification facility for instrument support or technical assistance. Work in the lab of Y.C. was supported by grants from the National Key R&D Program of China (2017YFA0503503), the CAS Pilot Strategic Science and Technology Projects B (XDB08030201), the NSFC (31861143028, 31670754, and 31872714), NSFC-ISF (31861143028), the CAS Major Science and Technology Infrastructure Open Research Projects, and the CAS-Shanghai Science Research Center (CAS-SSRC-YH-2015-01, DSS-WXJZ-2018-0002). Work in the lab of X.Z. was supported by grants from the National Key R&D Program of China (2017YFA0503503) and Chinese Academy of Sciences (XDB19020102). Z.D. was supported by a National Postdoctoral Program for Innovative Talents (BX201700262), NSFC (31800623), and the China Postdoctoral Science Foundation (2017M621550).

## Additional information

### Funding

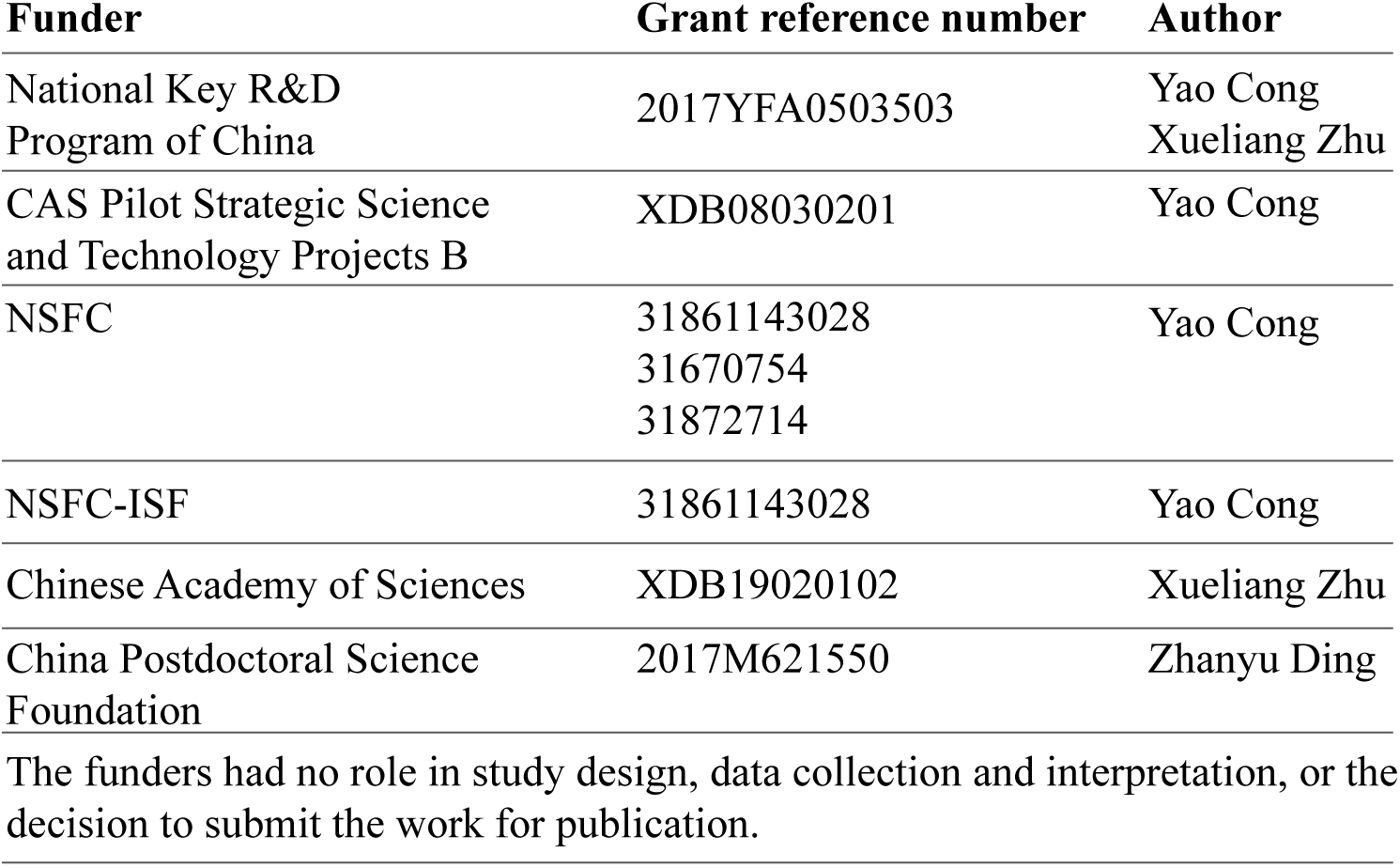

### Author contributions

Y.C., X.Z., and X.Y. designed the experiments, F.L., W.Z., and Q.G. purified proteins, F.L., H.L., and L.Z., performed functional analysis, W.Z. performed EM data collection, W.Z., Z.D., J.L., and Z.F. performed data analysis and modeling, C.P. performed the XL-MS experiments, W.Z., Y.C., Z.D, X.Z., and X.Y. analyzed the structure, W.Z., F.L. Y.C., X.Z., X.Y. and Z.D. wrote the manuscript.

### Data availability

All data needed to evaluate the conclusions in the paper are present in the paper and/or the Supplementary Materials. Cryo-EM map has been deposited in the EMDB with the accession number of ***. Additional data related to this paper may be requested from the authors.

## References

Abbasi F, Miyata H, Shimada K, Morohoshi A, Nozawa K, Matsumura T, Xu Z, Pratiwi P, and Ikawa M. 2018. RSPH6A is required for sperm flagellum formation and male fertility in mice. J Cell Sci 131. DOI: 10.1242/jcs.221648, PMID: 30185526

Baker ML, Abeysinghe SS, Schuh S, Coleman RA, Abrams A, Marsh MP, Hryc CF, Ruths T, Chiu W, and Ju T. 2011. Modeling protein structure at near atomic resolutions with Gorgon. Journal of structural biology 174:360–373. DOI: 10.1016/j.jsb.2011.01.015, PMID: 21296162

Baker ML, Ju T, and Chiu W. 2007. Identification of secondary structure elements in intermediate-resolution density maps. Structure 15:7–19. DOI: 10.1016/j.str.2006.11.008, PMID: 17223528

Barbato A, Frischer T, Kuehni CE, Snijders D, Azevedo I, Baktai G, Bartoloni L, Eber E, Escribano A, Haarman E, et al. 2009. Primary ciliary dyskinesia: a consensus statement on diagnostic and treatment approaches in children. Eur Respir J 34:1264–1276. DOI: 10.1183/09031936.00176608, PMID: 19948909

Brokaw CJ. 2009. Thinking about flagellar oscillation. Cell Motil Cytoskeleton 66:425–436. DOI: 10.1002/cm.20313, PMID: 18828155

Brokaw CJ, Luck DJ, and Huang B. 1982. Analysis of the movement of Chlamydomonas flagella:” the function of the radial-spoke system is revealed by comparison of wild-type and mutant flagella. J Cell Biol 92:722–732. DOI: 10.1083/jcb.92.3.722, PMID: 7085755

Bui KH, and Ishikawa T. 2013. 3D structural analysis of flagella/cilia by cryo-electron tomography. Methods Enzymol 524:305–323. DOI: 10.1016/b978-0-12-397945-2.00017-2, PMID: 23498747

Bui KH, Sakakibara H, Movassagh T, Oiwa K, and Ishikawa T. 2008. Molecular architecture of inner dynein arms in situ in Chlamydomonas reinhardtii flagella. J Cell Biol 183:923–932. DOI: 10.1083/jcb.200808050, PMID: 19029338

Bush A, Chodhari R, Collins N, Copeland F, Hall P, Harcourt J, Hariri M, Hogg C, Lucas J, Mitchison HM, et al. 2007. Primary ciliary dyskinesia: current state of the art. Archives of disease in childhood 92:1136–1140. DOI: 10.1136/adc.2006.096958, PMID: 17634184

Carbajal-Gonzalez BI, Heuser T, Fu X, Lin J, Smith BW, Mitchell DR, and Nicastro D. 2013. Conserved structural motifs in the central pair complex of eukaryotic flagella. Cytoskeleton (Hoboken) 70:101–120. DOI: 10.1002/cm.21094, PMID: 23281266

Castleman VH, Romio L, Chodhari R, Hirst RA, de Castro SC, Parker KA, Ybot-Gonzalez P, Emes RD, Wilson SW, Wallis C, et al. 2009. Mutations in radial spoke head protein genes RSPH9 and RSPH4A cause primary ciliary dyskinesia with centralmicrotubular-pair abnormalities. Am J Hum Genet 84:197–209. DOI: 10.1016/j.ajhg.2009.01.011, PMID: 19200523

Curry AM, Williams BD, and Rosenbaum JL. 1992. Sequence analysis reveals homology between two proteins of the flagellar radial spoke. Molecular and cellular biology 12:3967–3977. DOI: PMID: 1508197

Delgehyr N, Meunier A, Faucourt M, Bosch Grau M, Strehl L, Janke C, and Spassky N. 2015. Ependymal cell differentiation, from monociliated to multiciliated cells. Methods Cell Biol 127:19–35. DOI: 10.1016/bs.mcb.2015.01.004, PMID: 25837384

Diener DR, Yang P, Geimer S, Cole DG, Sale WS, and Rosenbaum JL. 2011. Sequential assembly of flagellar radial spokes. Cytoskeleton (Hoboken) 68:389–400. DOI: 10.1002/cm.20520, PMID: 21692193

Ding Z, Fu Z, Xu C, Wang Y, Wang Y, Li J, Kong L, Chen J, Li N, Zhang R, et al. 2017. High-resolution cryo-EM structure of the proteasome in complex with ADP-AlFx. Cell Res 27:373–385. DOI: 10.1038/cr.2017.12, PMID: 28106073

Ding Z, Xu C, Sahu I, Wang Y, Fu Z, Huang M, Wong CCL, Glickman MH, and Cong Y. 2019. Structural Snapshots of 26S Proteasome Reveal Tetraubiquitin-Induced Conformations. Mol Cell 73:1150–1161.e1156. DOI: 10.1016/j.molcel.2019.01.018, PMID: 30792173

Frommer A, Hjeij R, Loges NT, Edelbusch C, Jahnke C, Raidt J, Werner C, Wallmeier J, Grosse-Onnebrink J, Olbrich H, et al. 2015. Immunofluorescence Analysis and Diagnosis of Primary Ciliary Dyskinesia with Radial Spoke Defects. Am J Respir Cell Mol Biol 53:563–573. DOI: 10.1165/rcmb.2014-0483OC, PMID: 25789548

Gaillard AR, Diener DR, Rosenbaum JL, and Sale WS. 2001. Flagellar radial spoke protein 3 is an A-kinase anchoring protein (AKAP). J Cell Biol 153:443–448. DOI: 10.1083/jcb.153.2.443, PMID: 11309423

Goodenough UW, and Heuser JE. 1985. Substructure of inner dynein arms, radial spokes, and the central pair/projection complex of cilia and flagella. J Cell Biol 100:2008–2018. DOI: 10.1083/jcb.100.6.2008, PMID: 2860115

Grant T, Rohou A, and Grigorieff N. 2018. cisTEM, user-friendly software for singleparticle image processing. Elife 7. DOI: 10.7554/eLife.35383, PMID: 29513216

Hirokawa N, Tanaka Y, Okada Y, and Takeda S. 2006. Nodal flow and the generation of left-right asymmetry. Cell 125:33–45. DOI: 10.1016/j.cell.2006.03.002, PMID: 16615888

Huang B, Piperno G, Ramanis Z, and Luck DJ. 1981. Radial spokes of Chlamydomonas flagella: genetic analysis of assembly and function. J Cell Biol 88:80–88. DOI: 10.1083/jcb.88.1.80, PMID: 7204490

Ishikawa T. 2013. 3D structure of eukaryotic flagella/cilia by cryo-electron tomography. Biophysics (Nagoya-shi) 9:141–148. DOI: 10.2142/biophysics.9.141, PMID: 27493552

Jeanson L, Copin B, Papon JF, Dastot-Le Moal F, Duquesnoy P, Montantin G, Cadranel J, Corvol H, Coste A, Desir J, et al. 2015. RSPH3 Mutations Cause Primary Ciliary Dyskinesia with Central-Complex Defects and a Near Absence of Radial Spokes. Am J Hum Genet 97:153–162. DOI: 10.1016/j.ajhg.2015.05.004, PMID: 26073779

Jin M, Han W, Liu C, Zang Y, Li J, Wang F, Wang Y, and Cong Y. 2019. An ensemble of cryo-EM structures of TRiC reveal its conformational landscape and subunit specificity. Proceedings of the National Academy of Sciences of the United States of America 116:19513–19522. DOI: 10.1073/pnas.1903976116, PMID: 31492816

Jivan A, Earnest S, Juang YC, and Cobb MH. 2009. Radial spoke protein 3 is a mammalian protein kinase A-anchoring protein that binds ERK1/2. J Biol Chem 284:29437–29445. DOI: 10.1074/jbc.M109.048181, PMID: 19684019

Kelley LA, Mezulis S, Yates CM, Wass MN, and Sternberg MJ. 2015. The Phyre2 web portal for protein modeling, prediction and analysis. Nat Protoc 10:845–858. DOI: 10.1038/nprot.2015.053, PMID: 25950237

Kim DE, Chivian D, and Baker D. 2004. Protein structure prediction and analysis using the Robetta server. Nucleic Acids Res 32:W526–531. DOI: 10.1093/nar/gkh468, PMID: 15215442

Knowles MR, Daniels LA, Davis SD, Zariwala MA, and Leigh MW. 2013. Primary ciliary dyskinesia. Recent advances in diagnostics, genetics, and characterization of clinical disease. Am J Respir Crit Care Med 188:913–922. DOI: 10.1164/rccm.201301-0059CI, PMID: 23796196

Knowles MR, Ostrowski LE, Leigh MW, Sears PR, Davis SD, Wolf WE, Hazucha MJ, Carson JL, Olivier KN, Sagel SD, et al. 2014. Mutations in RSPH1 cause primary ciliary dyskinesia with a unique clinical and ciliary phenotype. Am J Respir Crit Care Med 189:707–717. DOI: 10.1164/rccm.201311-2047OC, PMID: 24568568

Kohno T, Wakabayashi K, Diener DR, Rosenbaum JL, and Kamiya R. 2011. Subunit interactions within the Chlamydomonas flagellar spokehead. Cytoskeleton (Hoboken) 68:237–246. DOI: 10.1002/cm.20507, PMID: 21391306

Kott E, Legendre M, Copin B, Papon JF, Dastot-Le Moal F, Montantin G, Duquesnoy P, Piterboth W, Amram D, Bassinet L, et al. 2013. Loss-of-function mutations in RSPH1 cause primary ciliary dyskinesia with central-complex and radial-spoke defects. Am J Hum Genet 93:561–570. DOI: 10.1016/j.ajhg.2013.07.013, PMID: 23993197

Kucukelbir A, Sigworth FJ, and Tagare HD. 2014. Quantifying the local resolution of cryo-EM density maps. Nat Methods 11:63–65. DOI: 10.1038/nmeth.2727, PMID: 24213166

Leigh MW, Pittman JE, Carson JL, Ferkol TW, Dell SD, Davis SD, Knowles MR, and Zariwala MA. 2009. Clinical and genetic aspects of primary ciliary dyskinesia/Kartagener syndrome. Genetics in medicine: official journal of the American College of Medical Genetics 11:473–487. DOI: 10.1097/GIM.0b013e3181a53562, PMID: 19606528

Lin J, Heuser T, Carbajal-Gonzalez BI, Song K, and Nicastro D. 2012. The structural heterogeneity of radial spokes in cilia and flagella is conserved. Cytoskeleton (Hoboken) 69:88–100. DOI: 10.1002/cm.21000, PMID: 22170736

Lin J, Yin W, Smith MC, Song K, Leigh MW, Zariwala MA, Knowles MR, Ostrowski LE, and Nicastro D. 2014. Cryo-electron tomography reveals ciliary defects underlying human RSPH1 primary ciliary dyskinesia. Nat Commun 5:5727. DOI: 10.1038/ncomms6727, PMID: 25473808

Luck D, Piperno G, Ramanis Z, and Huang B. 1977. Flagellar mutants of Chlamydomonas: studies of radial spoke-defective strains by dikaryon and revertant analysis. Proceedings of the National Academy of Sciences of the United States of America 74:3456–3460. DOI: PMID: 269405

Ludtke SJ, Baldwin PR, and Chiu W. 1999. EMAN: semiautomated software for high-resolution single-particle reconstructions. Journal of structural biology 128:82–97. DOI: 10.1006/jsbi.1999.4174, PMID: 10600563

Mitchell DR, and Nakatsugawa M. 2004. Bend propagation drives central pair rotation in Chlamydomonas reinhardtii flagella. J Cell Biol 166:709–715. DOI: 10.1083/jcb.200406148, PMID: 15337779

Nicastro D, Schwartz C, Pierson J, Gaudette R, Porter ME, and McIntosh JR. 2006. The molecular architecture of axonemes revealed by cryoelectron tomography. Science (New York, NY) 313:944–948. DOI: 10.1126/science.1128618, PMID: 16917055

Oda T, Yanagisawa H, Yagi T, and Kikkawa M. 2014. Mechanosignaling between central apparatus and radial spokes controls axonemal dynein activity. J Cell Biol 204:807–819. DOI: 10.1083/jcb.201312014, PMID: 24590175

Pigino G, Bui KH, Maheshwari A, Lupetti P, Diener D, and Ishikawa T. 2011. Cryoelectron tomography of radial spokes in cilia and flagella. J Cell Biol 195:673–687. DOI: 10.1083/jcb.201106125, PMID: 22065640

Piperno G, Huang B, Ramanis Z, and Luck DJ. 1981. Radial spokes of Chlamydomonas flagella: polypeptide composition and phosphorylation of stalk components. J Cell Biol 88:73–79. DOI: 10.1083/jcb.88.1.73, PMID: 6451632

Rohou A, and Grigorieff N. 2015. CTFFIND4: Fast and accurate defocus estimation from electron micrographs. Journal of structural biology 192:216–221. DOI: 10.1016/j.jsb.2015.08.008, PMID: 26278980

Rosenthal PB, and Henderson R. 2003. Optimal determination of particle orientation, absolute hand, and contrast loss in single-particle electron cryomicroscopy. J Mol Biol 333:721–745. DOI: 10.1016/j.jmb.2003.07.013, PMID: 14568533

Satir P, Heuser T, and Sale WS. 2014. A Structural Basis for How Motile Cilia Beat. Bioscience 64:1073–1083. DOI: 10.1093/biosci/biu180, PMID: 26955066

Shinohara K, Chen D, Nishida T, Misaki K, Yonemura S, and Hamada H. 2015. Absence of Radial Spokes in Mouse Node Cilia Is Required for Rotational Movement but Confers Ultrastructural Instability as a Trade-Off. Dev Cell 35:236–246. DOI: 10.1016/j.devcel.2015.10.001, PMID: 26506310

Smith EF, and Yang P. 2004. The radial spokes and central apparatus: mechanochemical transducers that regulate flagellar motility. Cell Motil Cytoskeleton 57:8–17. DOI: 10.1002/cm.10155, PMID: 14648553

Sturgess JM, Chao J, Wong J, Aspin N, and Turner JA. 1979. Cilia with defective radial spokes: a cause of human respiratory disease. N Engl J Med 300:53–56. DOI: 10.1056/nejm197901113000201, PMID: 152870

Tang G, Peng L, Baldwin PR, Mann DS, Jiang W, Rees I, and Ludtke SJ. 2007. EMAN2: an extensible image processing suite for electron microscopy. Journal of structural biology 157:38–46. DOI: 10.1016/j.jsb.2006.05.009, PMID: 16859925

van Heel M, and Schatz M. 2005. Fourier shell correlation threshold criteria. Journal of structural biology 151:250–262. DOI: 10.1016/j.jsb.2005.05.009, PMID: 16125414

Wang H, Han W, Takagi J, and Cong Y. 2018a. Yeast Inner-Subunit PA-NZ-1 Labeling Strategy for Accurate Subunit Identification in a Macromolecular Complex through Cryo-EM Analysis. J Mol Biol 430:1417–1425. DOI: 10.1016/j.jmb.2018.03.026, PMID: 29625202

Wang Y, Ding Z, Liu X, Bao Y, Huang M, Wong CCL, Hong X, and Cong Y. 2018b. Architecture and subunit arrangement of the complete Saccharomyces cerevisiae COMPASS complex. Sci Rep 8:17405. DOI: 10.1038/s41598-018-35609-8, PMID: 30479350

Wargo MJ, and Smith EF. 2003. Asymmetry of the central apparatus defines the location of active microtubule sliding in Chlamydomonas flagella. Proceedings of the National Academy of Sciences of the United States of America 100:137–142. DOI: 10.1073/pnas.0135800100, PMID: 12518061

Warner FD. 1970. New observations on flagellar fine structure. The relationship between matrix structure and the microtubule component of the axoneme. J Cell Biol 47:159–182. DOI: 10.1083/jcb.47.1.159, PMID: 4935335

Warner FD, and Satir P. 1974. The structural basis of ciliary bend formation. Radial spoke positional changes accompanying microtubule sliding. J Cell Biol 63:35–63. DOI: 10.1083/jcb.63.1.35, PMID: 4424314

Wirschell M, Zhao F, Yang C, Yang P, Diener D, Gaillard A, Rosenbaum JL, and Sale WS. 2008. Building a radial spoke: flagellar radial spoke protein 3 (RSP3) is a dimer. Cell Motil Cytoskeleton 65:238–248. DOI: 10.1002/cm.20257, PMID: 18157907

Witman GB, Plummer J, and Sander G. 1978. Chlamydomonas flagellar mutants lacking radial spokes and central tubules. Structure, composition, and function of specific axonemal components. J Cell Biol 76:729–747. DOI: 10.1083/jcb.76.3.729, PMID: 632325

Yang B, Wu YJ, Zhu M, Fan SB, Lin J, Zhang K, Li S, Chi H, Li YX, Chen HF, et al. 2012. Identification of cross-linked peptides from complex samples. Nat Methods 9:904–906. DOI: 10.1038/nmeth.2099, PMID: 22772728

Yang P, Diener DR, Yang C, Kohno T, Pazour GJ, Dienes JM, Agrin NS, King SM, Sale WS, Kamiya R, et al. 2006. Radial spoke proteins of Chlamydomonas flagella. J Cell Sci 119:1165–1174. DOI: 10.1242/jcs.02811, PMID: 16507594

Yang P, Yang C, and Sale WS. 2004. Flagellar radial spoke protein 2 is a calmodulin binding protein required for motility in Chlamydomonas reinhardtii. Eukaryot Cell 3:72–81. DOI: 10.1128/ec.3.1.72-81.2004, PMID: 14871938

Zang Y, Jin M, Wang H, Cui Z, Kong L, Liu C, and Cong Y. 2016. Staggered ATP binding mechanism of eukaryotic chaperonin TRiC (CCT) revealed through high-resolution cryo-EM. Nat Struct Mol Biol 23:1083–1091. DOI: 10.1038/nsmb.3309, PMID: 27775711

Zariwala MA, Knowles MR, and Omran H. 2007. Genetic defects in ciliary structure and function. Annu Rev Physiol 69:423-450. DOI: 10.1146/annurev.physiol.69.040705.141301, PMID: 17059358

Zhao H, Zhu L, Zhu Y, Cao J, Li S, Huang Q, Xu T, Huang X, Yan X, and Zhu X. 2013. The Cep63 paralogue Deup1 enables massive de novo centriole biogenesis for vertebrate multiciliogenesis. Nat Cell Biol 15:1434–1444. DOI: 10.1038/ncb2880 ncb2880 [pii], PMID: 24240477

Zheng SQ, Palovcak E, Armache JP, Verba KA, Cheng Y, and Agard DA. 2017. MotionCor2: anisotropic correction of beam-induced motion for improved cryoelectron microscopy. Nat Methods 14:331–332. DOI: 10.1038/nmeth.4193, PMID: 28250466

Zhu L, Liu H, Chen Y, Yan X, and Zhu X. 2019. Rsph9 is critical for ciliary radial spoke assembly and central pair microtubule stability. Biol Cell 111:29–38. DOI: 10.1111/boc.201800060, PMID: 30383886

Zivanov J, Nakane T, Forsberg BO, Kimanius D, Hagen WJ, Lindahl E, and Scheres SH. 2018. New tools for automated high-resolution cryo-EM structure determination in RELION-3. Elife 7. DOI: 10.7554/eLife.42166, PMID: 30412051

