## supplemental materials for "Distinct architecture and composition of mouse axonemal radial spoke head revealed by cryo-EM"

### Supplementary Information

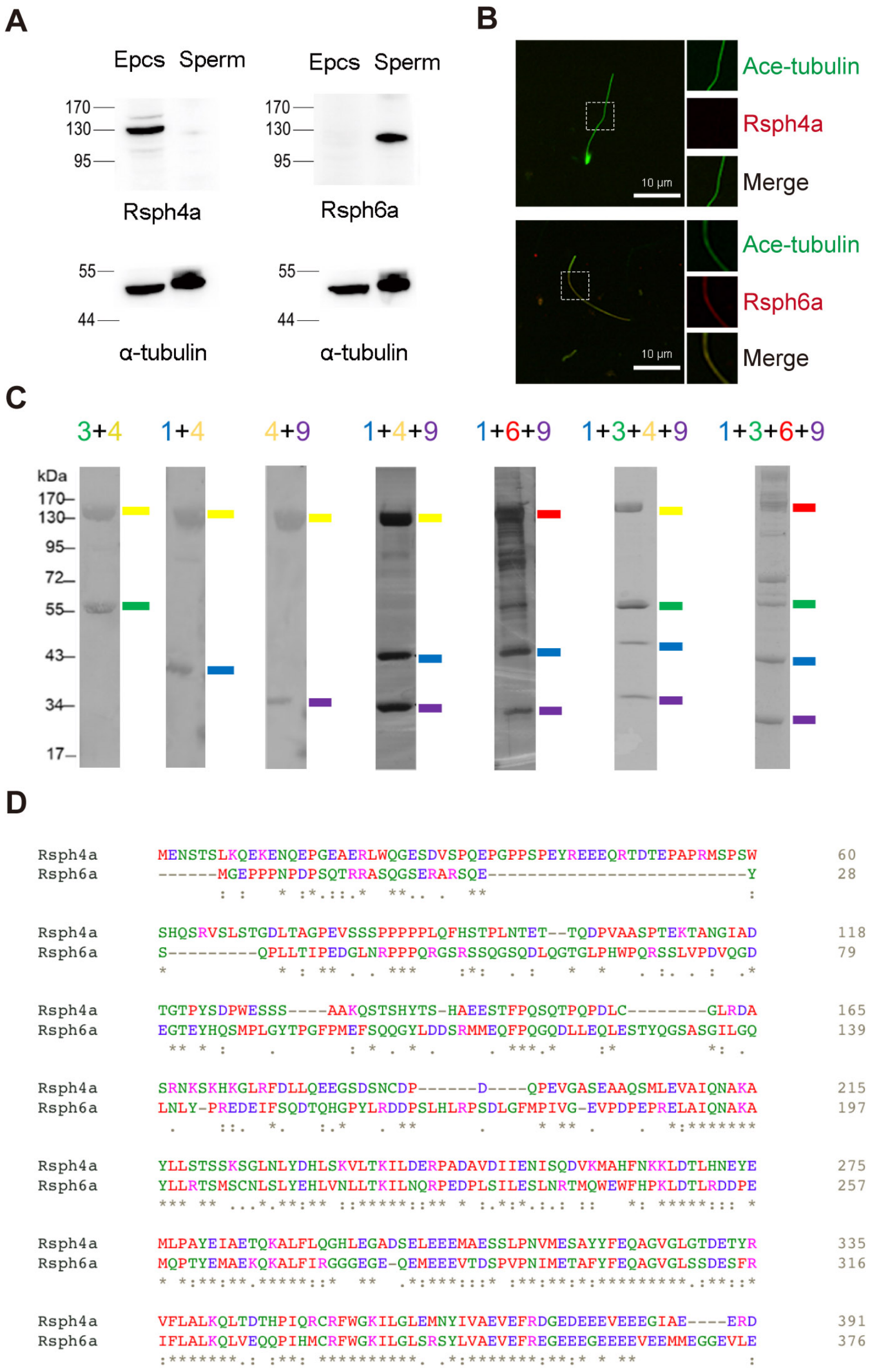


**Figure 1—figure supplement 1. Comparisons between Rsph4a and Rsph6a and assembly process of RS head core complex. (A, B)** Rsph4a and Rsph6a expressions are tissue specific. **(A)** Western blot experiments in mEPCs and sperm lysate showed Rsph4a specifically expressed in mouse mEPCs while Rsph6a specifically expressed in mouse

Figure S1 continued on next page

matured sperm. **(B)** Immunofluorescent of mouse matured sperm confirmed that Rsph6a specifically expressed in mouse sperm. Ace-tubulin served as a flagella axoneme marker. **(C)** Coomassie-blue staining, Ponceau's staining or silver staining SDS-PAGE of binary and multimeric RS head complexes purified by Flag tandem affinity purification. The RS head proteins are indicated with color-coded marks. Variances in gel mobility of the individual components is caused by using different protein as bait. **(D)** Sequence similarity between Rsph4a and Rsph6a by Clustal Omega. Asterisk marks indicates positions which have a single, fully conserved residue, period marks indicate the conservation between groups of weakly similar properties, colon marks indicate conservation between groups of strongly similar properties.

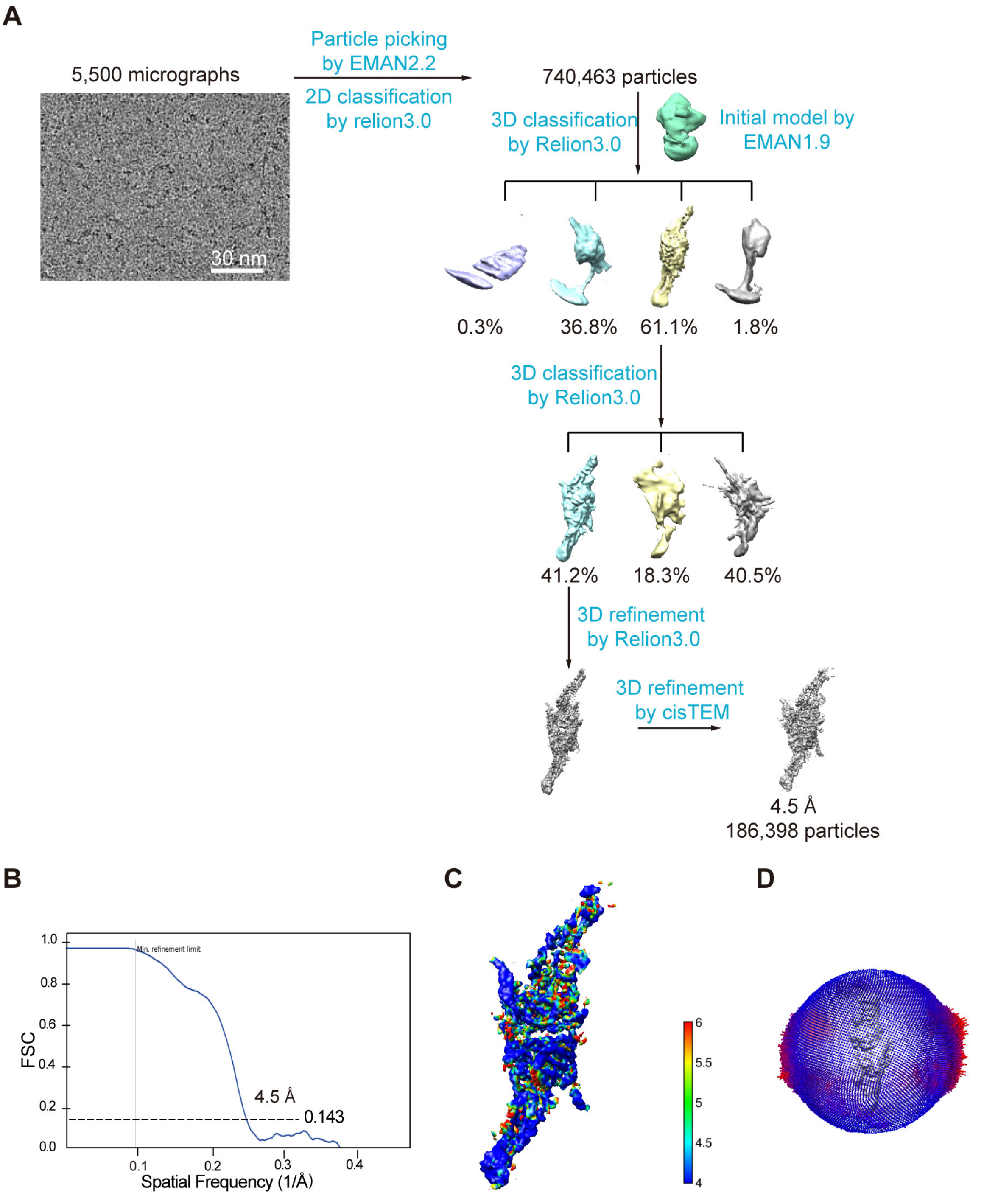


**Figure 2—figure supplement 1. Work flow for the cryo-EM data processing of RS head core complex. (A)** 3D classification and refinement procedures. After multiple rounds of 3D classification, we obtained a 3D map of RS head core complex. Details are described in the Methods. **(B)** Resolution estimation of the cryo-EM map of the RS head

Figure S2 continued on next page

core complex according to the gold-standard FSC criterion of 0.143. **(C)** Local resolution estimation of the core complex by Resmap. **(D)** The angular distribution of the 3D reconstruction, indicating unequal occupancy of angular classes.

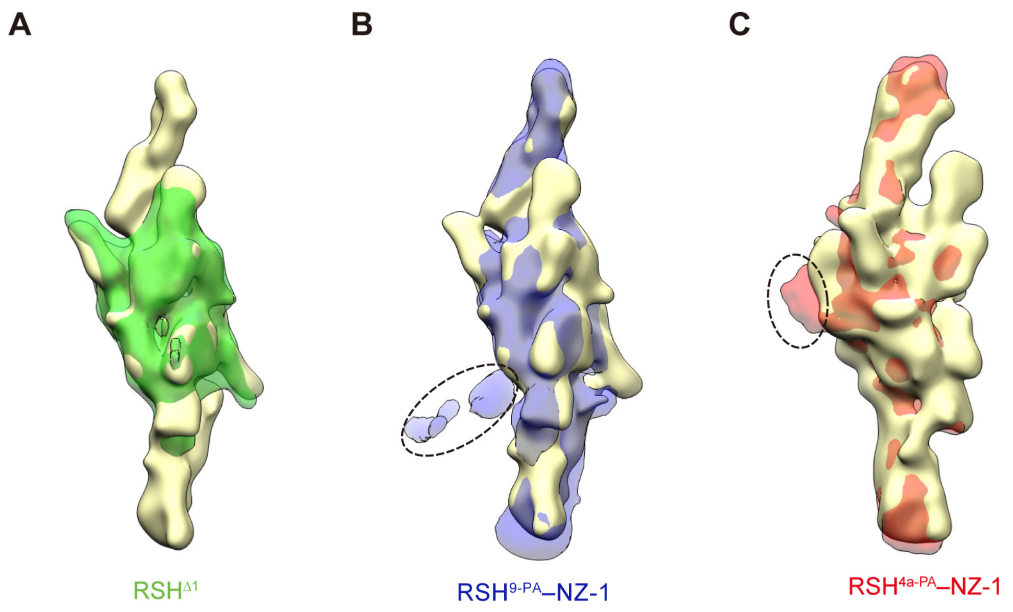


**Figure 3—figure supplement 1. Identification of RS head subunits determined through subunit deletion and PA-NZ-1 epitope labeling strategies. (A)** Alignment of the RSH$^{\Delta 1}$ map (green transparent) with a blurred RS head core map in similar 15 Å resolution (in yellow surface throughout this figure). The two arms are disappeared compared with the WT core complex. Showing is the front view. **(B)** Alignment of the RSH$^{9\text{-}PA}$–NZ-1 map (blue transparent) with the blurred WT core complex. A patch of exposed extra density corresponding to the NZ-1 Fab is indicated by a black ellipsoid. **(C)** Alignment of the RSH$^{4a\text{-}PA}$–NZ-1 map (red transparent) with the blurred RS head core map. An exposed extra density corresponding to NZ-1 Fab is indicated by black ellipsoid. Showing is the side view.

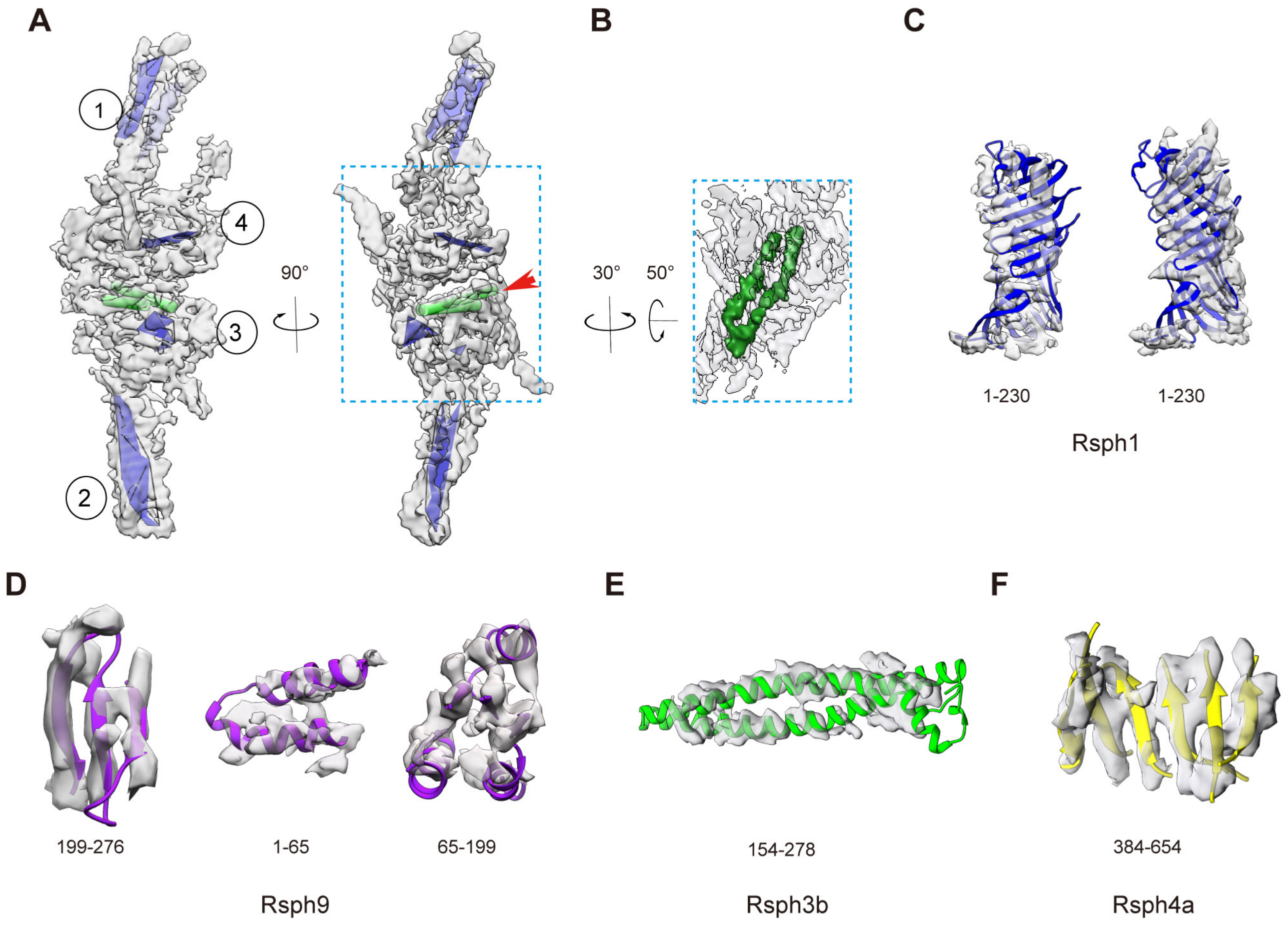


**Figure 3—figure supplement 2. Structural analysis of RS head core complex. (A)** Typical SSEs (β-sheet feature indicated by blue surface and α-helix feature by green cylinder) of our RS head core map determined by Gorgon analysis. Here we used ①, ②, ③, and ④ to indicate the locations of detectable β-sheets, and the red arrow to indicate the location of a two-long-helix bundle. **(B)** Visualization of the two-long-helix bundle (in green) in the RS head map (transparent grey), with a rotation relative to the right panel in **(A)** for better visualization. **(C-F)** Reasonably good matching between model and map especially for the Gorgon determined typical SSE regions, e.g. we can visualize the β-strand separation in Rsph9 **(D)** and Rsph4a **(F)** subunits.

**A**

Swiss models

| Subunit | Length (aa) | Template structure | Sequence coverage | Sequence identify（%） |
|---|---|---|---|---|
| Rsph1 | 301 | 1n6c | 64-174 （36.8%） | 23.73 |
| Rsph3b | 389 | 4xa4 | 109-245 （35.2%） | 16 |
| | | 5jxd | 240-323（21.6%） | 17.74 |
| Rsph4a | 716 | 4f9k | 211-282 （10.1%） | 6 |
| Rsph9 | 276 | 3o02 | 5-32（10.1%） | 27.59 |

Robetta models

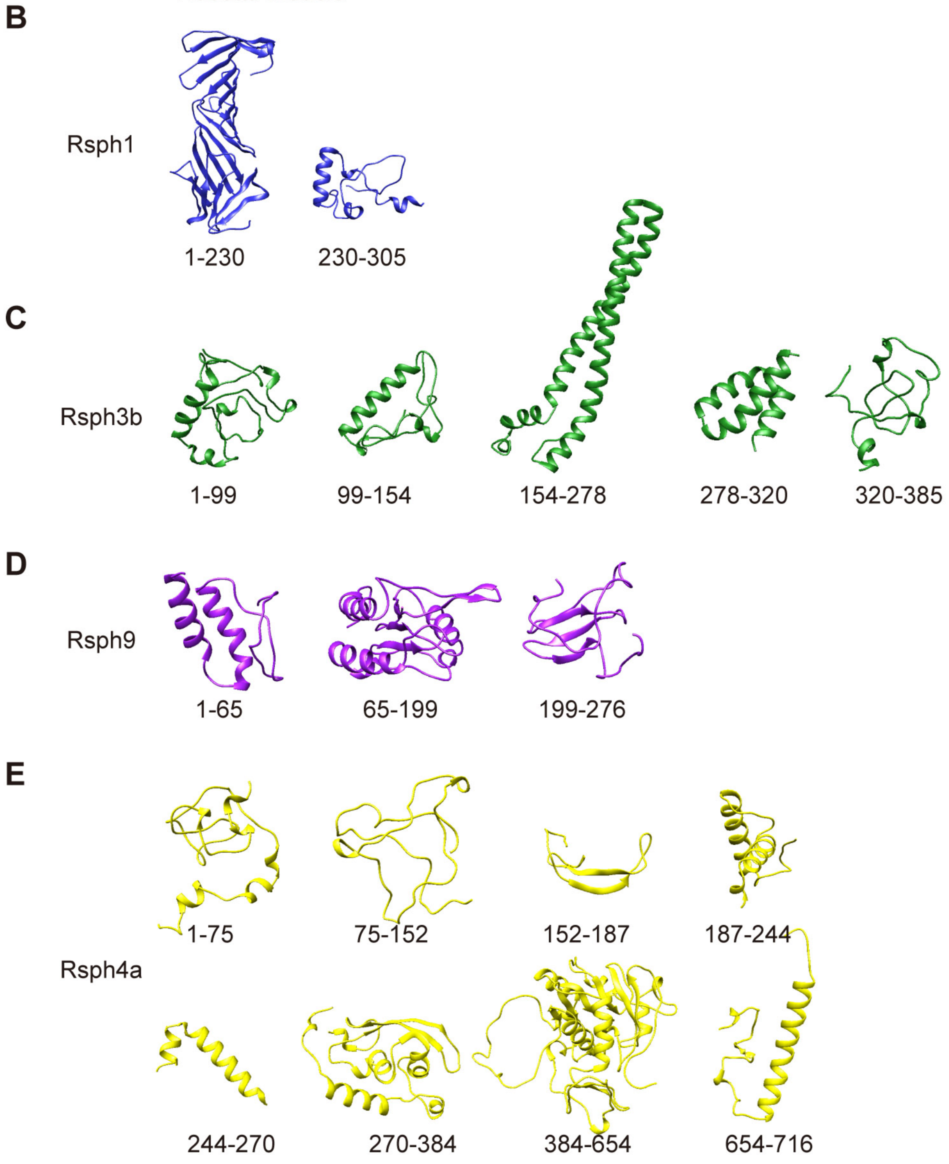

**Figure 3—figure supplement 3. Model building for RS head core complex. (A)** Template model detected through SWISS Model server. **(B-E)** Comparative and de-novo models of individual subunits predicted by Robetta server. The models have the aa numbers labeled underneath.

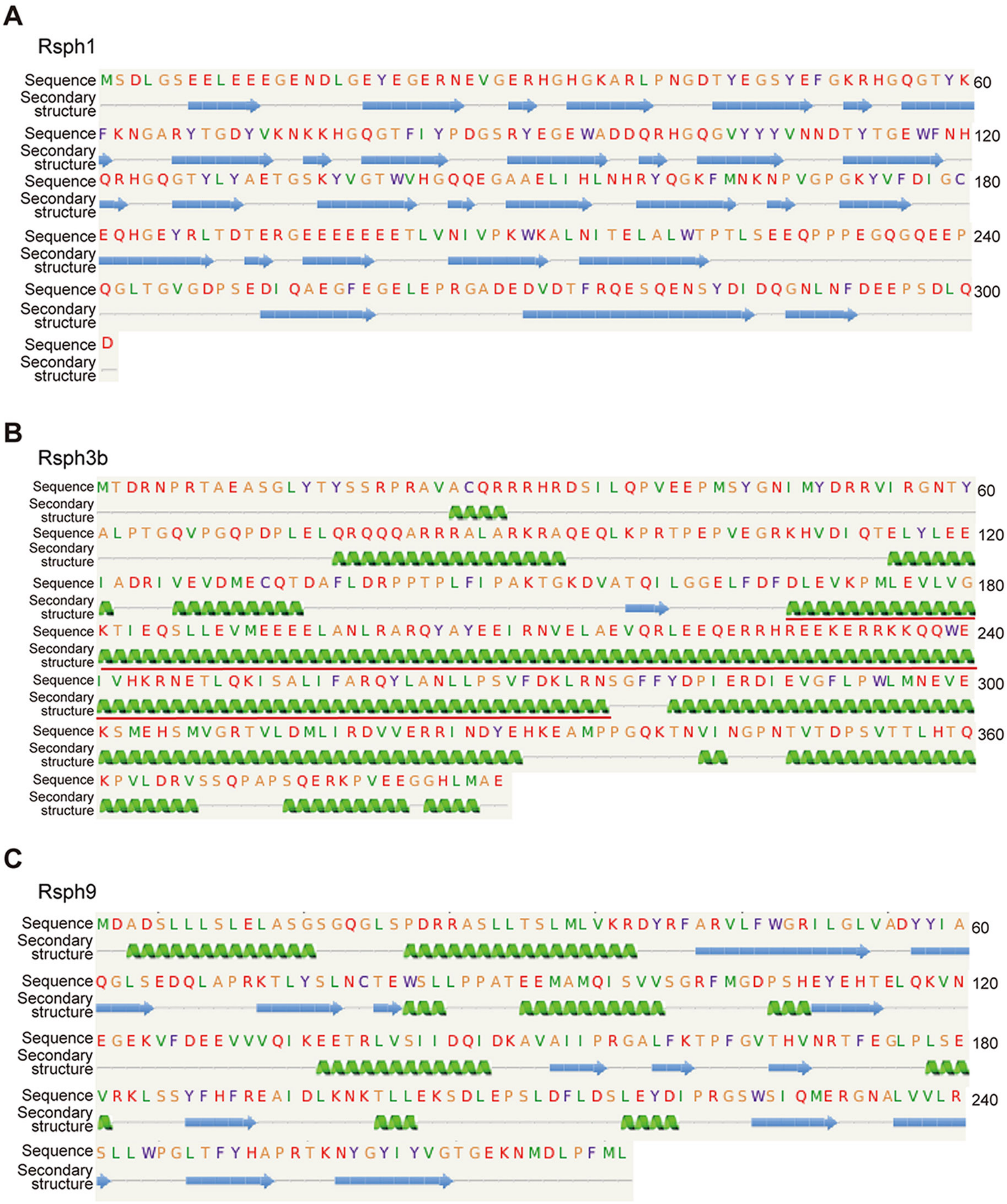
A
Rsph1
Sequence
Secondary structure
60
120
180
240
300
B
Rsph3b
Sequence
Secondary structure
60
120
180
240
300
360
C
Rsph9
Sequence
Secondary structure
60
120
180
240

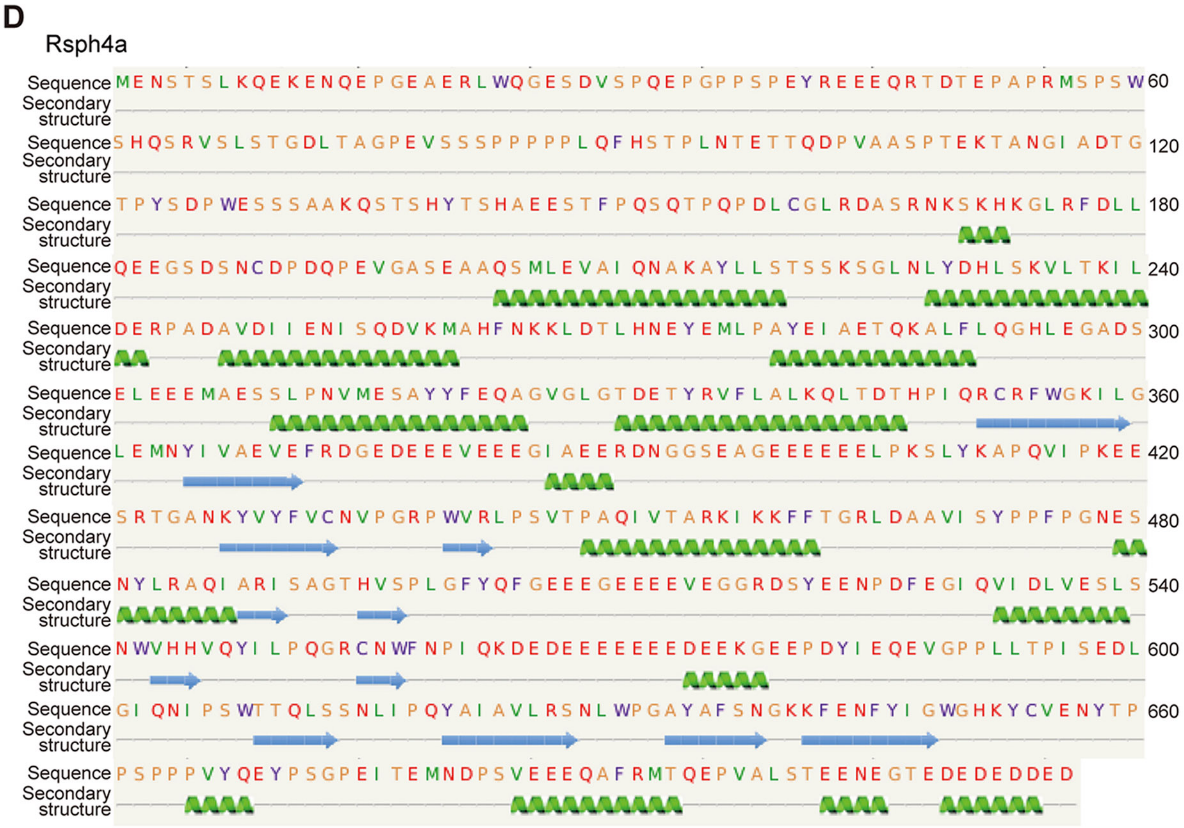


**Figure 3—figure supplement 4. Subunit secondary structure prediction of the RS head core complex by Phyre2. (A-D)** SSE prediction for Rsph1 **(A)**, Rsph3b **(B)**, Rsph9 **(C)**, and Rsph4a **(D)**. Green ribbon represents α-helix, and blue arrow represents β-strand. For Rsph3b, the two extremely long α-helices (~154 aa to 278 aa) were indicated by red underline **(B)**.

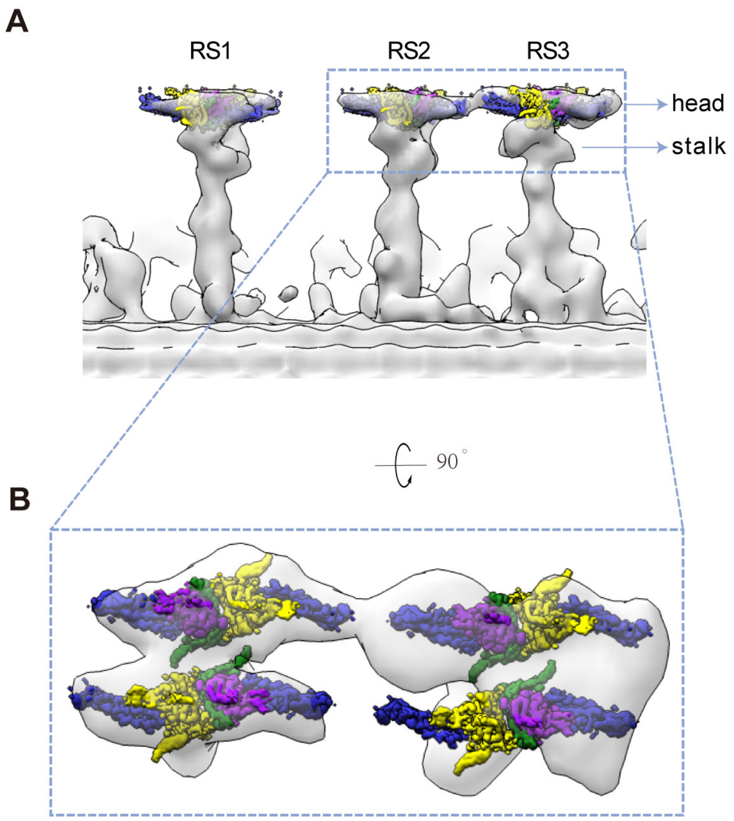


**Figure 4—figure supplement 1. Fitting of our RS head core map into the cryo-ET map of WT human DMT. (A)** Longitudinal (proximal end of the axoneme to the left) view of RSs from the cryo-ET map of WT human axoneme doublet microtubule (transparent grey, EMD-5950) fitted with our cryo-EM map of RS head core complex (density in color). **(B)** Magnified top view of RS2 and RS3 fitted with our RS head core complex, indicating that unlike RS2, RS3 cannot hold the RS head core map very well.

```
RSP3            ----MVQAKAQQQLYTHAAEPKAVQQRRAKYRE---DETTQTLPTANIMFDRRVVRGNTY      53
Rsph3b          MTDRNPRTAEASGLYTYSSRPRAVACQRRRHRDSILQPVEEPMSYGNIMYDRRVIRGNTY      60
                      ::   . ***:::.*:**  :* ::*:   : . : :  .***:****:*****

RSP3            AARILPADA------TQTQTKGPSPASTKKRTTRTLPPRTPEAVDGRRHIDIQTDVYLEE      107
Rsph3b          ALPTGQVPGQPDPLELQRQQQARRRALARKRAQEQLKPRTPEPVEGRKHVDIQTELYLEE      120
                *     . .       * * :.   * ::**: . * ***** *:**:*:****::****

RSP3            LTDTVPEADTSTQTDAFLDRPPTPLFVPQKTGTDAITQIENGDLFDFDFEVEPILEVLVG      167
Rsph3b          IADRIVEVDMECQTDAFLDRPPTPLFIPAKTGKDVATQILGGELFDFDLEVKPMLEVLVG      180
                ::* : *.* . **************:* ***.*. *** .*:*****:**:*:******

RSP3            KVLEQGLMEVLEEEELAAMRAHQEHFEQIRNAELVATQRMEAAERRKLEEKERRMQQERE      227
Rsph3b          KTIEQSLLEVMEEEELANLRARQYAYEEIRNVELAEVQRLEEQERRHREEKERRKKQQWE      240
                *.:**.*:**:****** :**:*  :*:***.**. .**:*  ***: ****** :*: *

RSP3            RVERERVVRQKVAASAFARGYLSGIVNTVFDRLVSSGYIYDPVMREVETAFMPWLKEQAI      287
Rsph3b          IVHKRNETLQKISALIFARQYLANLLPSVFDKLRNSGFFYDPIERDIEVGFLPWLMNEVE      300
                 *.:.. . **::*  *** **:.:: :***:* .**::***: *::*..*:*** ::.

RSP3            GYLARGVVARRVVDKLVEDAAAALAANRSTLADKAASTAATVDAWAERQAKMEAELQGKE      347
Rsph3b          KSMEHSMVGRTVLDMLIRDVVERRINDYEH---KEAMPPGQ----------KTNVINGPN      347
                  : :.:*.* *:* *:.*..     : .    * *   .               ::* :

RSP3            LEAVRRRPTFVLRELKPAVASADAVEAAAAELTAQAEEAANAKWEADKAEAAEKARAEAE      407
Rsph3b          T--VTDPSVTTLHTQKPVLDRVSSQPAPSQER-KPV------------------------      380
                   *    . .*:  **.:  ..:  * : *    .

RSP3            AAAEEQKALLEELAATAAAEAEERGEEPPAEPPSLPDGVEPVDVEAEVAKAVEAVPKPPV      467
Rsph3b          ---EEGGHLMAE------------------------------------------------      389
                   **   *: *

RSP3            KEVTDIDILSYMMDKGAITKDAIIQALAVHALGDKAYTNHPAFAEAEGA   516
Rsph3b          -------------------------------------------------   389
```

**Figure 4—figure supplement 2. Sequence alignment between *Chlamydomonas* RSP3 and mouse Rsph3b.**

**Table S1. Cryo-EM data collection and refinement statistics**

| | |
|---|---|
| EM equipment Voltage (kV) | Titan Krios 300 |
| Detector | K2 Summit |
| Pixel size (Å) | 1.02 |
| Total electron dose ($e^-$/Å$^2$) | 58 |
| Dose rate ($e^-$/physical pixel/sec) | 8 |
| Exposure time (s) | 7.6 |
| Defocus range (μm) | -1.2 ~ -2.2 |
| Reconstruction software | Relion3.0, cisTEM |
| Original particles | 740,043 |
| Final particles | 186,398 |
| Symmetry | C1 |
| Final resolution (Å) | 4.5 |

**Table S2. XL-MS detected subunit interactions of RS head core complex**

| Protein1(site)-protein2(site) | Peptides | Best e-value | Spec count |
|---|---|---|---|
| R1(60)-R9(199) | HGQGTYKFK(7)-NKTLLEK(2) | 4.11E-03 | 1 |
| R1(62)-R4(459) | FKNGAR(2)-KFFTGR(1) | 3.84E-06 | 1 |
| R1(73)-R4(418) | YTGDYVKNK(7)-APQVIPKEESR(7) | 1.75E-06 | 1 |
| R1(73)-R4(459) | YTGDYVKNK(7)-KFFTGR(1) | 1.11E-06 | 3 |
| R1(75)-R3(231) | NKKHGQGTFIYPDGSR(2)-HREEKER(5) | 6.55E-02 | 1 |
| R1(76)-R4(459) | HGQGTFIYPDGSR(1)-KFFTGR(1) | 2.39E-11 | 3 |
| R1(76)-R4(411) | KHGQGTFIYPDGSR(1)-SLYKAPQVIPK(4) | 4.45E-07 | 2 |
| R1(162)-R3(97) | YQGKFMNK(4)-AQEQLKPR(6) | 1.42E-03 | 1 |
| R1(162)-R4(266) | YQGKFMNK(4)-KLDTLHNEYEMLPAYEIAETQK(1) | 3.68E-02 | 1 |
| R1(162)-R4(459) | YQGKFMNK(4)-KFFTGR(1) | 1.47E-09 | 7 |
| R1(166)-R4(459) | FMNKNPVGPGK(4)-KFFTGR(1) | 4.61E-04 | 1 |
| R4(223)-R9(197) | AYLLSTSSKSGLNLYDHLSK(9)-EAIDLKNK(6) | 1.79E-02 | 1 |
| R4(234)-R3(97) | SGLNLYDHLSKVLTK(11)-AQEQLKPR(6) | 3.08E-04 | 2 |
| R4(266)-R3(97) | KLDTLHNEYEMLPAYEIAETQK(1)-AQEQLKPR(6) | 1.87E-02 | 2 |
| R4(341)-R3(97) | VFLALKQLTDTHPIQR(6)-AQEQLKPR(6) | 2.64E-02 | 2 |
| R4(411)-R3(97) | SLYKAPQVIPK(4)-AQEQLKPR(6) | 4.57E-04 | 5 |
| R4(411)-R3(231) | SLYKAPQVIPK(4)-HREEKER(5) | 1.79E-06 | 2 |
| R4(411)-R3(236) | SLYKAPQVIPK(4)-KQQWEIVHK(1) | 6.00E-06 | 1 |
| R4(411)-R9(199) | SLYKAPQVIPK(4)-NKTLLEK(2) | 1.60E-08 | 2 |
| R4(418)-R3(97) | APQVIPKEESR(7)-AQEQLKPR(6) | 5.99E-06 | 4 |
| R4(418)-R3(231) | APQVIPKEESR(7)-HREEKER(5) | 2.41E-08 | 2 |
| R4(418)-R9(197) | APQVIPKEESR(7)-EAIDLKNK(6) | 2.81E-03 | 1 |
| R4(456)-R3(231) | LPSVTPAQIVTARKIK(14)-EEKERR(3) | 7.19E-02 | 2 |
| R4(459)-R3(97) | KFFTGR(1)-AQEQLKPR(6) | 5.06E-05 | 4 |
| R4(459)-R3(231) | KFFTGR(1)-HREEKER(5) | 7.02E-02 | 1 |
| R4(459)-R3(244) | QQWEIVHKR(8)-KFFTGR(1) | 6.16E-04 | 1 |
| R4(459)-R9(199) | KFFTGR(1)-NKTLLEK(2) | 1.24E-08 | 5 |
| R4(459)-R9(256) | KFFTGR(1)-TKNYGYIYVGTGEK(2) | 1.37E-04 | 1 |
| R9(161)-R3(97) | GALFKTPFGVTHVNR(5)-AQEQLKPR(6) | 9.92E-03 | 1 |
| R9(197)-R3(236) | EAIDLKNK(6)-KQQWEIVHK(1) | 1.49E-02 | 1 |
| R9(197)-R3(244) | EAIDLKNK(6)-QQWEIVHKR(8) | 3.97E-03 | 1 |
| R9(197)-R3(231) | EAIDLKNK(6)-HREEKER(5) | 3.34E-06 | 1 |
| R9(199)-R3(231) | NKTLLEK(2)-HREEKER(5) | 4.32E-02 | 1 |

Note: R1 indicates Rsph1, R3 indicates Rsph3b, R4 indicates Rsph4a, R9 indicates Rsph9.

#### Table S3. List of antibodies

| Primary antibodies | | | | | |
|---|---|---|---|---|---|
| **Antigen** | **Species** | **Supplier** | **Cat. #** | **IB** | **IF** |
| Ace-tub | mouse | Sigma-Aldrich | T6793 (6-11B-1) | 1:5000 | 1:1000 |
| Gapdh | rabbit | Proteintech Group Inc. | 10494-1-AP | 1:5000 | |
| HA | mouse | Sigma-Aldrich | H9658 | 1:5000 | |
| Flag | mouse | Sigma-Aldrich | F3165 | 1:5000 | |
| Rsph1 | rabbit | home-made | | 1:1000 | 1:500 |
| Rsph3b | rabbit | Proteintech | 17603-1-AP | 1:1000 | 1:500 |
| Rsph4a | rabbit | home-made | | 1:2000 | 1:2000 |
| Rsph6a | rabbit | home-made | | 1:1000 | 1:500 |
| Rsph9 | rabbit | home-made | | 1:1000 | 1:500 |
| Rsph10b | rabbit | home-made | | 1:1000 | 1:500 |

| Secondary antibodies | | | | | |
|---|---|---|---|---|---|
| **Name** | **Species** | **Supplier** | **Cat. #** | **IB** | **IF** |
| anti-Mouse IgG (H+L)-HRP | goat | Life Technologies | G-21040 | 1:5000 | |
| anti-Rabbit IgG (H+L)-HRP | goat | Life Technologies | G-21234 | 1:5000 | |
| anti-Mouse IgG (H+L)-Alexa Fluor 488 | donkey | Life Technologies | A-21202 | | 1:1000 |
| anti-Rabbit IgG (H+L)-Alexa Fluor 488 | donkey | Life Technologies | A-21206 | | 1:1000 |
| anti-Chicken IgY-Alexa Fluor 488 | goat | Life Technologies | A-11039 | | 1:1000 |
| anti-Rat IgG (H+L)-Alexa Fluor 488 | donkey | Jackson ImmunoResearch | 712-545-153 | | 1:1000 |
| anti-Mouse IgG (H+L)-Cy3 | donkey | Jackson ImmunoResearch | 715-165-151 | | 1:1000 |
| anti-Rabbit IgG (H+L)-Cy3 | donkey | Jackson ImmunoResearch | 711-165-152 | | 1:1000 |
| anti-Rat IgG ( H+L)-Alexa Fluor 546 | goat | Life Technologies | A11081 | | 1:1000 |
| anti-Mouse IgG (H+L)-Alexa Fluor 647 | goat | Life Technologies | A-21236 | | 1:1000 |

**Movie 1.** Coordination of our RS head core cryo-EM map (in red) into the framework of DMT-CP based on previous cryo-ET studies on sea urchin CP (EMD-9385, in dark cyan) and human DMT (EMD-5950, in purple). Based on this fitting, we proposed a sawtooth model in RS-CP interaction.
